# Multimeric nanobodies from camelid engineered mice and llamas potently neutralize SARS-CoV-2 variants

**DOI:** 10.1101/2021.03.04.433768

**Authors:** Jianliang Xu, Kai Xu, Seolkyoung Jung, Andrea Conte, Jenna Lieberman, Frauke Muecksch, Julio Cesar Cetrulo Lorenzi, Solji Park, Zijun Wang, Lino Tessarollo, Tatsiana Bylund, Gwo-Yu Chuang, Adam S. Olia, Tyler Stephens, I-Ting Teng, Yaroslav Tsybovsky, Tongqing Zhou, Theodora Hatziioannou, Paul D. Bieniasz, Michel C. Nussenzweig, Peter D. Kwong, Rafael Casellas

## Abstract

Since the start of the coronavirus disease-2019 (COVID-19) pandemic, severe acute respiratory syndrome coronavirus-2 (SARS-CoV-2) has caused more than 2 million deaths worldwide. Multiple vaccines have been deployed to date, but the continual evolution of the viral receptor-binding domain (RBD) has recently challenged their efficacy. In particular, SARS-CoV-2 variants originating in the U.K. (B.1.1.7), South Africa (B.1.351) and New York (B.1.526) have reduced neutralization activity from convalescent sera and compromised the efficacy of antibody cocktails that received emergency use authorization. Whereas vaccines can be updated periodically to account for emerging variants, complementary strategies are urgently needed to avert viral escape. One potential alternative is the use of camelid VHHs (also known as nanobodies), which due to their small size can recognize protein crevices that are inaccessible to conventional antibodies. Here, we isolate anti-RBD nanobodies from llamas and “nanomice” we engineered to produce VHHs cloned from alpacas, dromedaries and camels. Through binding assays and cryo-electron microscopy, we identified two sets of highly neutralizing nanobodies. The first group expresses VHHs that circumvent RBD antigenic drift by recognizing a region outside the ACE2-binding site that is conserved in coronaviruses but is not typically targeted by monoclonal antibodies. The second group is almost exclusively focused to the RBD-ACE2 interface and fails to neutralize pseudoviruses carrying the E484K or N501Y substitutions. Notably however, they do neutralize the RBD variants when expressed as homotrimers, rivaling the most potent antibodies produced to date against SARS-CoV-2. These findings demonstrate that multivalent nanobodies overcome SARS-CoV-2 variant mutations through two separate mechanisms: enhanced avidity for the ACE2 binding domain, and recognition of conserved epitopes largely inaccessible to human antibodies. Therefore, while new SARS-CoV-2 mutants will continue to emerge, nanobodies represent promising tools to prevent COVID-19 mortality when vaccines are compromised.

## Generation and characterization of nanomice

In contrast to mouse and human antibodies which are ∼150 kDa in size, the variable domain of camelids heavy chain-only antibodies (VHHs, also referred to as nanobodies or Nbs) retain full antigen specificity at a molecular weight of ∼15 kDa. Their smaller size and extended CDRs allow Nbs to bind epitopes not normally accessible to conventional antibodies, such as deep crevices and concave surfaces. Additional advantages of Nbs include higher expression yields, enhanced solubility, and tissue penetration. These features make Nbs ideal therapeutics, particularly against human viruses which often mask conserved epitopes with glycan shields. Despite their advantages, Nbs are still not widely used. One reason is that camelids are large animals that are expensive to maintain and not suitable for academic facilities. In addition, the lack of species inbreeding results in highly heterogenous immune responses. In a search for alternatives, naïve (non-immune) Nb libraries have been developed, where large numbers of VHHs with randomized CDR3s are synthesized and screened *in vitro* by phage or yeast display (McMahon et al., 2018; Moutel et al., 2016). However, in the absence of antibody affinity maturation in germinal centers *in vivo* (Victora and Nussenzweig, 2012), high specificity and diversity are difficult to achieve by this approach.

To bypass these hurdles, we sought to produce Nbs in mice by combining 18 alpaca, 7 dromedary and 5 camel VHH genes in a 25Kb insertion cassette (Figure 1A). Each gene was fused to a functional mouse VH promoter, leader exons, and a downstream recombination signal sequence (RSS) to ensure physiological expression and recombination (Figure S1). By means of CRISPR-Cas9 and recombinase-mediated cassette exchange (RMCE), the VHH cassette was inserted in lieu of the entire VH locus in mouse ES cells (Figure 1A).

**Figure 1:**
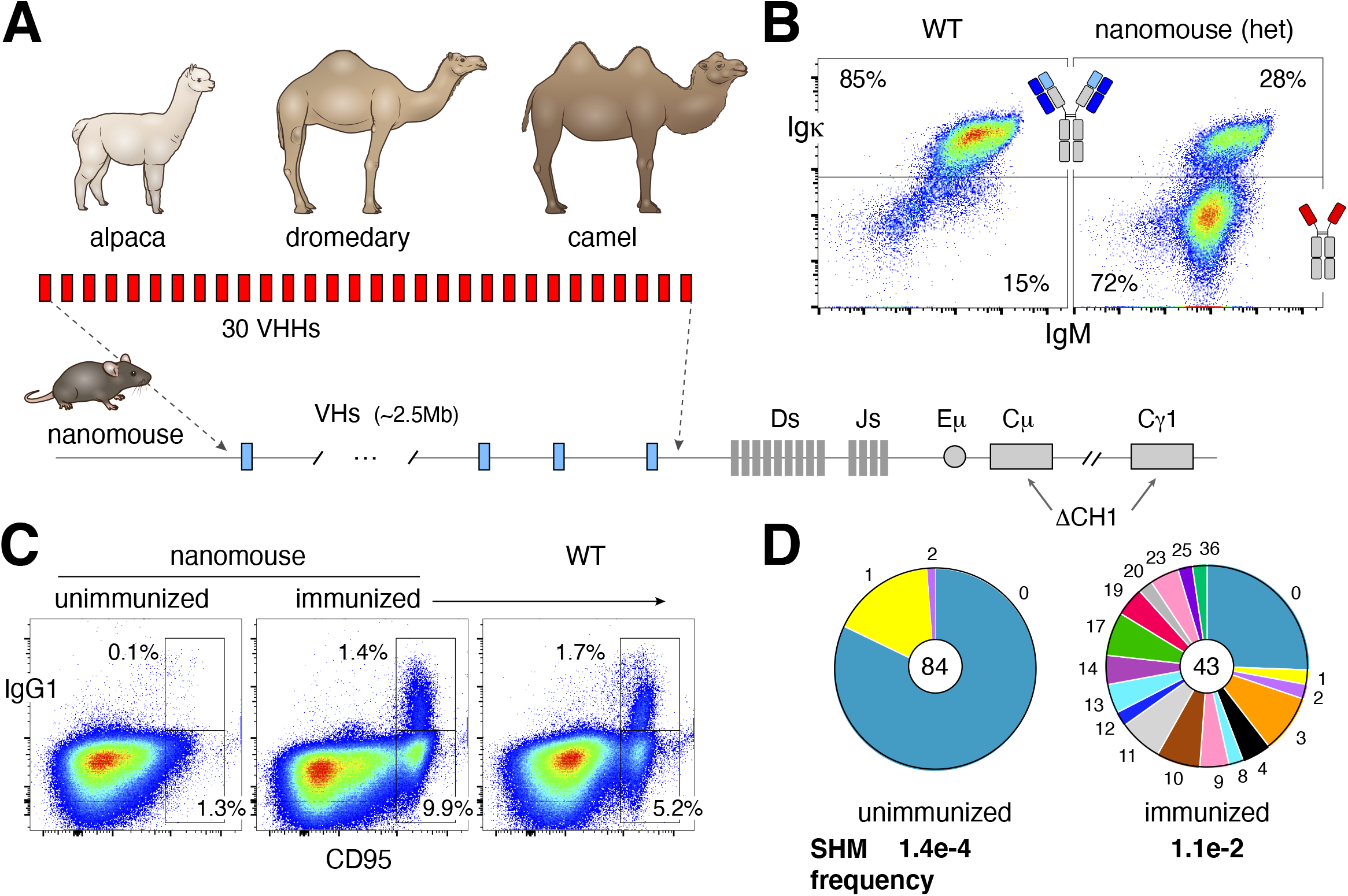
Production and characterization of nanomice. (**A**) 30 VHHs were selected from alpaca, dromedary and camel to create a cassette, which was inserted via CRISPR-Cas9 in lieu of the mouse VH locus (2.5MB in size). The CH1 exon of C µ and Cγ1 was also deleted to avoid misfolding of the antibody heavy chain when unable to pair with light chains. (**B**) Flow cytometry analysis of splenic B220+ B cells from WT or heterozygous nanomice. IgM+Igκ+ represent cells expressing conventional heavy-light chain antibodies, whereas IgM+Igκ-are mostly Igλ+ in WT (not shown) or single-chain antibody B cells in nanomice. (**C**) Flow cytometry analysis of splenic cells from unimmunized and immunized nanomice and controls stained with the germinal center markers CD95 and IgG1. (**D**) Pie charts showing VHH somatic hypermutation in unimmunized and immunized nanomice. Pie segments are proportional to the VHH sequences carrying the mutations indicated on the periphery of the chart. The middle circle contains the total number of sequences and the mutation frequency is shown in bold below.

Camelid Nbs are only expressed in conjunction with dedicated IgG2 and IgG3 that lack the CH1 domain (Muyldermans, 2013). In conventional antibodies, the hydrophobic surface of CH1 helps to pair heavy and light chain constant domains, while in dedicated camelid IgGs the CH1 exon is spliced out during transcription (Achour et al., 2008). To recapitulate this configuration in the mouse genome, we deleted CH1 from *Igγ1* in the VHH-targeted chromosome (Figure 1A). In addition, the *Igμ* CH1 was knocked out to prevent an arrest in B cell development, as previously reported in Nb-transgenic mice(Janssens et al., 2006). Mouse chimeras were derived from ES cells and the targeted allele was germline transmitted to F1 offspring (hereafter referred to as nanomice).

As expected, in wild type (WT) mice ∼85% of splenic B220^+^ B cells were IgM^+^Igκ^+^ (Figure 1B, left panel). In stark contrast, in heterozygous nanomice 72% of B lymphocytes displayed an IgM^+^Igκ^-^ phenotype (Figure 1B, right panel). Of these, less than 2% were IgM^+^Igλ^+^ (Figure S2A), implying that a large fraction of nanomouse B cells develop expressing single-chain antibodies. We confirmed this observation by amplifying VHH-DJ joining events using gene-specific primers. We found that all 30 VHHs were recombined to downstream JHs in bone marrow and spleen samples (Figure S2B).

For a more accurate quantification of VHH usage, VHH-DJ genes from three nanomouse spleens were cloned and sequenced. Analogous to the repertoire distribution of mouse and human VH genes (Demaison et al., 1995; Rettig et al., 2018), VHH usage varied considerably in nanomice, from <0.1% for VHH18 to 24% for VHH16 (Figure S2C). Importantly, VHH frequencies were reproducible in the three animals. We conclude that all VHH genes in nanomice undergo V(D)J recombination and are thus potentially available for expansion during the adaptive immune response.

Next, we generated VHH homozygous mice (*Igh*^*VHH/VHH*^) and characterized B cell development by flow cytometry. By and large, the B cell compartment of nanomice was normal, displaying all developmental stages including B1, B2 and marginal zone B cells (Figure S3A). One noticeable difference was an increased number of IgM^+^ transitional and immature B cells in the bone marrow and spleen, respectively. This is indicative of enhanced selection as cells transition from the short-to the long-lived B cell compartments. Consistent with this idea, CD23^high^CD21^low^ mature follicular B cells were reduced 1.7-fold in nanomice (Figure S3A). Another distinct feature was the absence of cell surface IgD (Figure S3B). This phenotype likely results from differential mRNA splicing due to CH1 deletion at *Igµ*, because IgD was also absent in *Igµ*-CH1^-/-^ only mice, where VHs and *Igγ1* CH1 are intact (Figure S3B). We conclude that mouse B cells can mature expressing single-chain antibodies.

## B cell activation and affinity maturation in nanomice

To probe B cell activation, splenic cells were isolated and cultured *ex-vivo* in the presence of lipopolysaccharide (LPS), interleukin-4 (IL-4) and αCD180 antibodies for 72-96h. Under these conditions, VHH-expressing cells underwent proliferation and class switch recombination to IgG1 (Figure S3B-C). To examine activation *in vivo*, we performed intraperitoneal immunizations with keyhole limpet hemocyanin (KLH) in complete Freund’s adjuvant. Twelve days post-immunization, nanomice showed equivalent numbers of B220^+^CD95^high^IgG1^+^ germinal center B cells relative to controls (Figure 1C).

To study affinity maturation against a specific antigen, we immunized nanomice with HIV-1 envelop trimer (BG505 DS-SOSIP), a component of HIV-1 entry machinery and an important target for HIV-1 vaccine efforts ((Kong et al., 2019), Figure S3D). Hypermutation of VHH genes in nanomice was increased relative to unimmunized controls (1.1e-2 vs. 1.4e-4, Figure 1D). The mutation spectra revealed an enrichment in G to A and C to T transitions (Figure S3E), consistent with the expected profile of activation-induced cytidine deaminase (AID) catalysis (Pham et al., 2003).

To measure the antibody response against BG505 DS-SOSIP, we characterized 16 nanobodies carrying different VHHs (1, 2, 6, 9, 16 and 19), which were enriched for HIV-1 trimer recognition. Sequence analysis showed CDR3s to be highly diverse in this group in terms of JH gene usage, mutations, and size (9-16 amino acids, Figure S4A). To measure binding kinetics, we next used bio-layer interferometry (BLI). The analysis identified four variants, all derived from VHH9, which displayed dissociation constants (*K*_D_s) ranging from 2 to 13 nM, demonstrating that they represent high-affinity binders (Figure S4B). Taken together, the data show that mouse B cells expressing single-chain antibodies can undergo efficient affinity maturation and produce highly specific Nbs upon immunization.

## Isolation of SARS-CoV-2 neutralizing nanobodies

We next sought to produce neutralizing Nbs against SARS-CoV-2. To this end, 3 nanomice and 1 llama were subjected to an 11-week immunization regimen which combined injections of recombinant receptor binding domain (RBD) and the stabilized prefusion spike (Figure 2A). Peripheral blood mononuclear cells (PBMCs) were isolated post-immunization and VHHs were amplified and cloned into a phagemid vector. Following one to two rounds of phage display, Nbs specific for the RBD were enriched using an ELISA-based binding screen. Sanger and deep-sequencing analyses before and after enrichment identified on average 26,000 Nb variants per library, representing a total of 192 unique CDR3s for llama and 199 for nanomice (Figure S5A). Nbs were then clustered by CDR3s and representative clones from each subgroup were isolated and tested for blocking RBD binding to the angiotensin-converting enzyme 2 (ACE2) receptor *in vitro* (Tan et al., 2020). Six llama and six nanomouse Nbs were selected with this method (Figure S5B-C).

**Figure 2:**
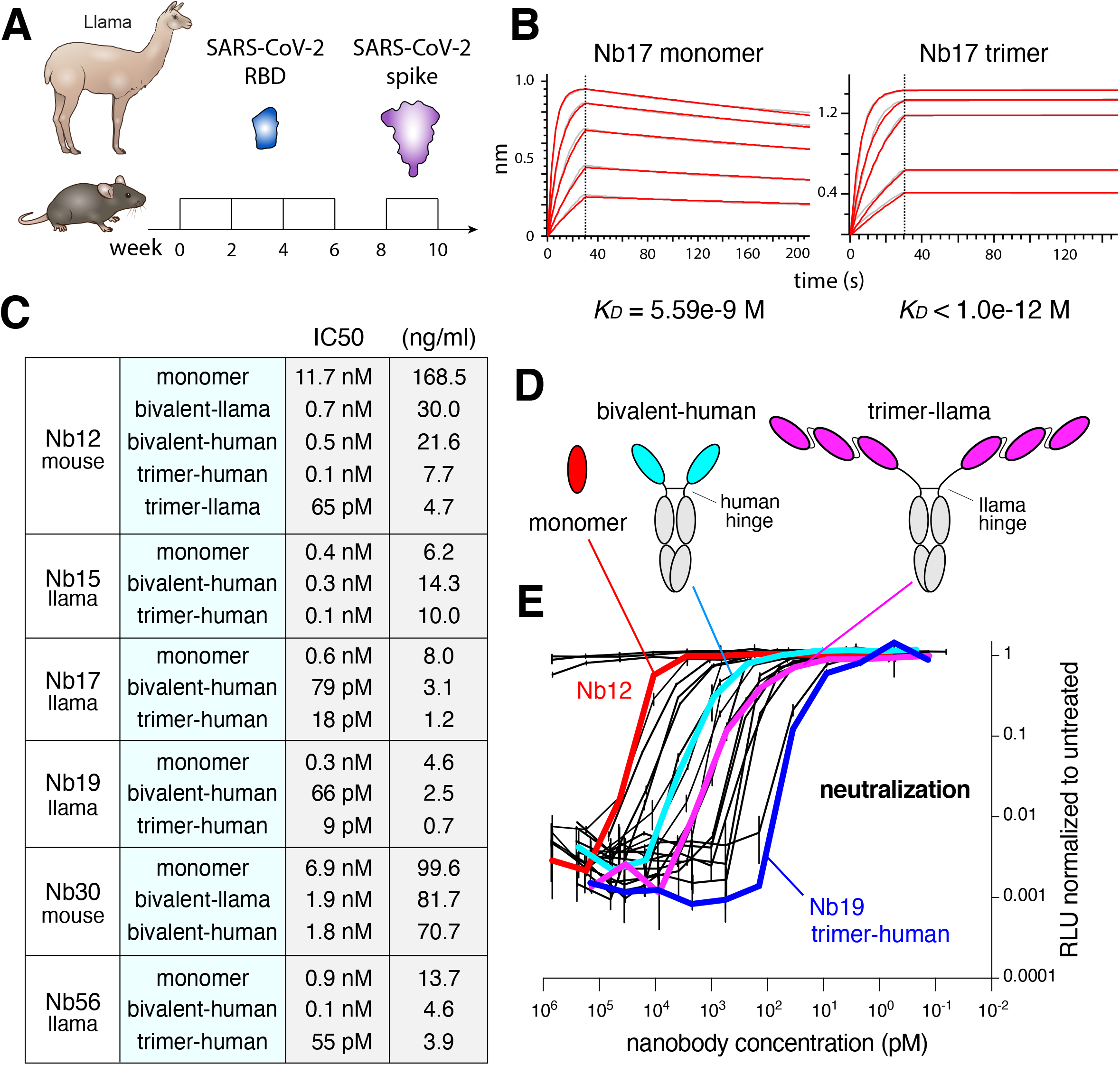
Isolation of Nbs against SARS-CoV-2. (**A**) Immunization regimen to obtain high affinity Nbs against the SARS-CoV-2 RBD from llama and nanomice. (**B**) BLI analysis of difference concentrations of Nb17 monomer and trimer binding to immobilized RBD. Red trace represents the raw data, and the kinetic fit is shown in grey underneath. Equilibrium (*KD*) constants are provided. (**C**) Table summarizing pseudovirus neutralization potency (IC50) of selected Nbs. Values are provided in molarity or as ng/ml. (**D**) Diagrams showing Nbs used in neutralization assays as monomers, bivalent or trimers (the last two fused to human IgG Fc via the human or llama hinge domain. (**E**) Neutralization of SARS-CoV-2 pseudovirus (luciferase assay) by the 20 Nb variants shown in panel **C**. Nb12 as monomer (red), pair (cyan) and trimer (magenta), as well as Nb19 as trimer (blue) are highlighted.

To refine the list of candidates we measured binding affinity for RBD by BLI. The analysis identified 4 llama (15, 17, 19, and 56) and 2 nanomouse (12 and 30) Nbs with dissociation constants tighter than 30 nM, indicating very high affinity for RBD (Figure 2B, S5D). The off-rate varied from 7.1×10^−3^ to 1.1×10^−3^ s^-1^, demonstrating slow dissociation for all Nbs (Figure S5D). We next explored neutralization *in vitro* using lentiviral particles pseudotyped with the SARS-CoV-2 spike (Robbiani et al., 2020). The Nb monomers displayed nM and sub-nM half-maximal inhibitory concentration (IC_50_), ranging from 11.7 nM (168.5 ng/ml) for Nb12 to 0.335 nM (4.6 ng/ml) for Nb19 (Figure 2C).

A crucial advantage of Nbs over conventional antibodies is that they can more easily be assembled into multimeric constructs, often resulting in remarkable avidity (Laursen et al., 2018; Schoof et al., 2020; Xiang et al., 2020). To explore this property, Nbs were fused as trimers using flexible GGGGS(x3) linkers of 15 amino acids in length. These constructs were connected to the human IgG1 Fc antibody portion, either via the human hinge domain or its much longer and flexible llama counterpart (Figure 2D). By fusing two VHH monomers to an IgG1 Fc we also created bivalent antibodies (Figure 2D). We found that neutralization increased with the number of linked monomers, from 3-fold for Nb15 to 180-fold for Nb12 (Figure 2C and 2E). Notably, the four most potent multimeric Nbs (12, 17, 19, and 56) reached IC50 values in the picomolar range, from 65 pM (4.7 ng/ml) to as little as 9 pM (0.7 ng/ml, Figure 2C and 2E). These values rank among the best reported to date for anti-SARS-CoV-2 Nbs (Saelens and Schepens, 2021).

## Multimeric nanobodies efficiently overcome SARS-CoV-2 escape mutants

With the worldwide spread of SARS-CoV-2, several variants carrying RBD mutations have emerged that increase transmissibility or allow escape from antibody neutralization. Of particular interest is the B.1.1.17 variant, containing an N501Y substitution, that has caused a recent upsurge in COVID-19 cases in the United Kingdom due to an estimated 70% increased transmissibility (Control, 2020). A second variant of concern is the South African B.1.351, which combines N501Y with two additional RBD substitutions: K417N and E484K. A third variant that has spread very rapidly in the New York area, B.1.526, shares the E484K mutation with the South African variant (Annavajhala et al., 2021; West et al., 2021). All of these mutations were shown to reduce the efficacy of serum antibodies elicited by the Moderna and Pfizer vaccines (Annavajhala et al., 2021; Wang et al., 2021a; Wang et al., 2021c; Wu et al., 2021).

We explored whether our leading Nbs could neutralize SARS-CoV-2 S pseudotyped viruses carrying the RBD variant mutations. The R683G substitution, which increases infectivity of pseudotyped viruses in vitro (Schmidt et al., 2020) was also included as a control. In contrast to their efficacy against the WT virus, Nb17, Nb19, and Nb56 were unable to neutralize viruses carrying the E484K mutation alone or in combination with K417N and N501Y (KEN construct, Figure 3A). Similarly, Nb15 was ineffective against the N501Y mutation. Surprisingly, however, with the exception of Nb17, they all remained highly potent neutralizers as trimers (Figure 3A-B). In the case of Nb15 and Nb56 trimers, IC50 values reached 30 pM and 14 pM respectively. Thus, the E484K and N501Y mutations allow viral escape from monomeric but not multimeric Nbs.

**Figure 3:**
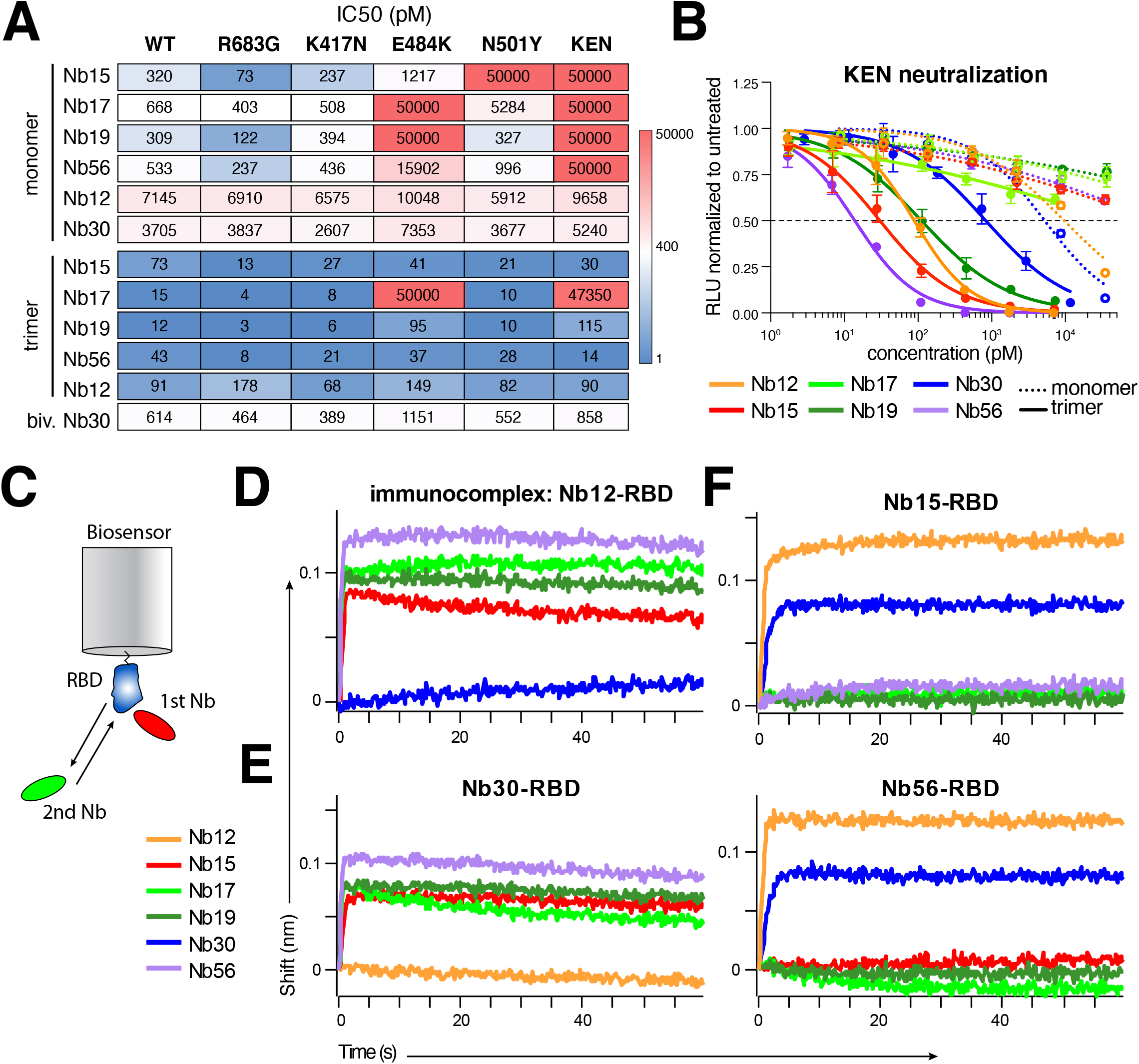
Neutralization of SARS-CoV-2 RBD WT and mutant pseudoviruses. (**A**) IC50 values for the selected Nbs based on neutralization assays of pseudoviruses carrying WT and the indicated mutant SARS-CoV-2 spike. Color gradient indicates values ranging from 0 (blue) to 50,000 pM (red). Pseudotyped viruses containing the E484K mutation (E484K and KEN) also contain the R683G mutation (for details see methods). (**B**) Neutralization assays, showing the sensitivity of pseudotyped viruses containing K417N/E484K/N501Y (KEN) SARS-CoV-2 spike proteins to isolated Nbs. Dotted lines represent activity of monomers whereas full lines represent that of trimers fused to human IgG1 Fc. Nb30 is bivalent. (**C**) Schematics summarizing BLI competition assay, where an Nb-RBD immunocomplex attached to the biosensor is incubated with a different Nb to measure binding. (**D-F**) Binding of Nbs to Nb12-RBD, Nb30-RBD, Nb15-RBD and Nb56-RBD immunocomplexes respectively.

In contrast to llama Nbs, nanomouse Nb12 and Nb30 were intriguing in that their neutralization potencies were unaltered by RBD mutations, either alone or in the KEN combination (Figure 3A-B). The strong implication is that llama and nanomouse Nbs recognize different regions on the RBD. To directly test this idea, we performed BLI experiments in which a preformed monomeric Nb-RBD immunocomplex was incubated with a second Nb (Figure 3C). We found that all four llama Nbs but not Nb30 could bind the Nb12-RBD immunocomplex (Figure 3D). Likewise, Nb30-RBD interfered with Nb12 binding, whereas llama Nbs could bind freely (Figure 3E). At the same time, Nb12 and Nb30 recognized all combinations of llama Nb-RBD complexes, whereas llama Nbs could not (Figure 3F and S6A). Thus, nanomouse and llama Nbs recognize two distinct neutralizing regions on SARS-CoV-2 RBD.

## Structural basis for Nb inhibition of SARS-CoV-2 WT and variants

To define the region bound by nanomouse Nbs, we collected single particle cryo-EM data on a Titian Krios for Nb12 and Nb30 in complex with HexaPro (Hsieh et al., 2020), a prefusion construct of the SARS-CoV-2 spike (Figure S7 and S8). In both cases, we used particle subtraction, classification and location refinement to enhance the resolution of the Nb-spike interface.

The structure of the Nb12-spike complex revealed Nb12 to induce a 2-RBD-up, 1 RBD-down spike conformation, with Nb12 recognizing a region towards the middle of the RBD, outside of the ACE2-binding region and distal from the 417, 484 and 501 mutations in emerging variants of concern (Figure 4A,4C and S9). Meanwhile, the structure of the Nb30-spike complex revealed Nb30 to induce a 3-RBD-up spike conformation, with Nb30 recognizing a region at the opposite end of RBD from the ACE2 binding motif and escape mutations (Figure 4B-C and S9).

**Figure 4:**
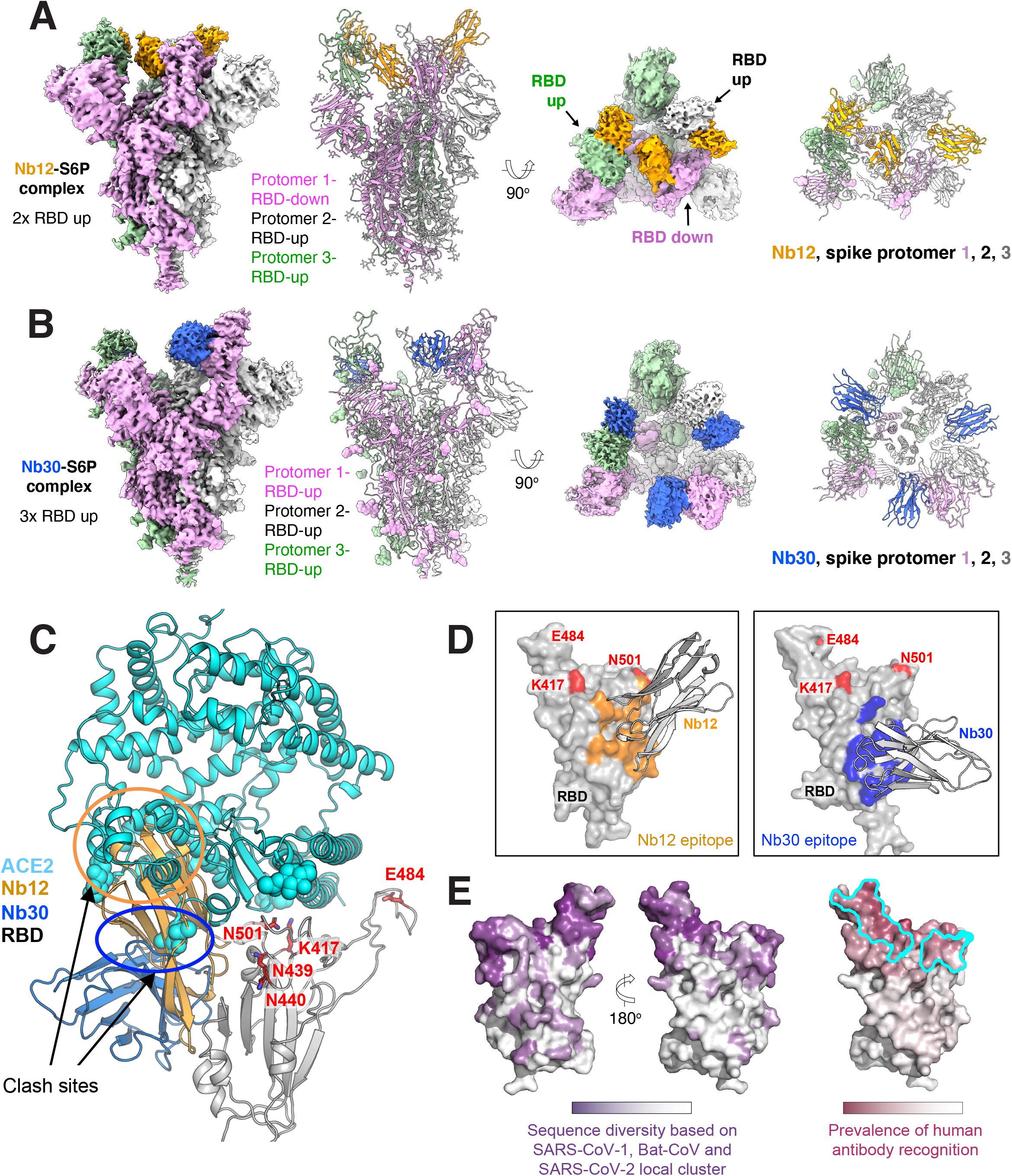
Structures of Nbs in complex with SARS-CoV-2 spike reveal nanomouse Nbs to recognize a conserved region, outside the ACE2-binding site. Cryo-EM structure of nanomouse Nb12 in complex with SARS-CoV-2 spike. (**B**) Cryo-EM structure of nanomouse Nb30 in complex with SARS-CoV-2 spike. (**C**) Cryo-EM defined structures of nanomouse Nbs recognize regions on RBD distal from emerging mutations 417, 484 and 501. (**D**) Interface between Nbs and spike. (**E**) Surface properties of RBD including sequence diversity (dark purple indicates high diversity), and prevalence of RBD-recognized regions by human antibodies (dark raspberry indicates high prevalence) and binding site for ACE2 (scribed in cyan).

To understand how the two nanomouse Nbs neutralized despite recognizing surfaces outside the ACE2 binding domain, we superimposed the structure of the ACE2-RBD complex (Benton et al., 2020; Lan et al., 2020; Zhou et al., 2020) with those of Nb12 and Nb30 (Figure 4C). We observed a substantial portion of Nb12 domain clashing with ACE2, indicating Nb12 and ACE2 binding to be sterically incompatible. With Nb30, a more subtle clash with glycan N322 on ACE2 was observed, which nonetheless also indicated Nb30 and ACE2 binding to be sterically incompatible.

To obtain a structural understanding of the neutralizing regions recognized by nanomouse and llama Nbs, we also determined 3d-negative stain EM reconstructions of each of the Nbs in complex with HexaPro spike. These reconstructions revealed the llama Nbs uniformly target the ACE2-binding interface, with Nb17, Nb19, and Nb56 inducing 1-RBD-up conformation, while Nb15 associates with all-RBD-down spikes (Figure S10). By contrast, both nanomouse Nb12 and Nb30 recognize RBD at a surface outside the ACE2-binding site (Figure 4D).

## RBD regions recognized by nanomouse versus human RBD-neutralizing antibodies

To provide insight into the prevalence of regions on RBD recognized by human antibodies versus nanomouse Nbs, we superimposed all of the RBD-directed human neutralizing antibodies in the PDB and quantified the recognition prevalence at the residue level (Figure 4E). While recognition extended over much of the RBD, the prevalence of human antibody recognition was much higher in the ACE2-binding region where emerging mutations reside (417, 484 and 501). By contrast, the regions recognized by Nb12 and Nb30 were more conserved and displayed substantially lower prevalence of human antibody recognition. Interestingly, the epitope of Nb12 overlaps considerably with those recognized by previous Nbs specific for SARS-CoV-1 (Wrapp et al., 2020) and -CoV-2 (Koenig et al., 2021; Xiang et al., 2020), raising the possibility that those Nbs might also block the new SARS-CoV-2 variants. The Nb30 binding footprint however is distanced further away from the ACE2 RBD motif and cover a surface area that is 79% conserved among the local cluster of coronaviruses including SARS-CoV-1, Bat-CoV and SARS-CoV-2, compared to 54% for Nb12 and on average 23% for human antibodies (Figure 4D-E). We conclude that the nanomouse VHHs circumvent RBD antigenic drift by recognizing a conserved region outside the ACE2-binding site.

## DISCUSSION

A key contribution of our study is the creation of Nb-producing mice. Previous work explored transgenic expression of 1-2 llama VHHs or human VHs in Igh^-/-^Igκ^-/-^Igλ^-/-^ mice (Drabek et al., 2016; Janssens et al., 2006; Teng et al., 2020). In our model, the 30 VHHs replace the entire VH domain leading to physiological recombination and selection during B cell development. While capable of producing high-affinity Nbs, our nanomouse 1.0 can be improved further by increasing the number of available VHH genes. This could be done by engineering a second allele carrying VHHs from llamas, vicuñas and guanacos, the three camelids not included in our model. We have identified >40 unique VHHs from these species, which when combined with the current allele should largely surpass the number of functional VHHs present in camelids (estimated to be ∼50 per species (Achour et al., 2008; Nguyen et al., 2000)). We anticipate that this and similar mouse models will help popularize the development of Nbs against infectious diseases or for basic applications. Just as important, the models may ultimately replace the use of camelids in biomedical research, thus protecting llamas and alpacas which are invariably culled in antibody production farms at the conclusion of a given study.

As a proof of principle, we used the nanomice to produce highly specific Nbs against SARS-CoV-2 RBD. Numerous monoclonal antibodies have been isolated so far from COVID-19 convalescent patients (Brouwer et al., 2020; Chi et al., 2020; Jones et al., 2020; Robbiani et al., 2020) and humanized mice immunized with the viral spike (Hansen et al., 2020). Notably, the most potent among them primarily block the RBD-ACE2 interface. Not surprisingly, immunotherapies involving such antibodies are highly vulnerable to escape variants carrying mutations at or around the ACE2 binding motif (Annavajhala et al., 2021; Wang et al., 2021a; Wang et al., 2021c; Wu et al., 2021). The mutations in some cases bestow higher affinity for ACE2 and appear to have been selected to evade the antibody response (Greaney et al., 2021; Wang et al., 2021b; Wibmer et al., 2021). To overcome the limitations of conventional antibodies we produced anti-RBD Nbs. As previous studies have shown, Nbs are well poised to accomplish this because of their versatility and capacity for multivalency (Esparza et al., 2020; Hanke et al., 2020; Huo et al., 2020; Koenig et al., 2021; Schoof et al., 2020; Tortorici et al., 2020; Wrapp et al., 2020).

From immunized llamas and nanomice we have isolated and characterized two sets of Nbs that effectively neutralize pseudotyped viruses carrying RBD mutations from U.K. and South African-New York variants. Similar to human antibodies, Nbs from the first group (Nb15, Nb56) hinder ACE2 binding to the spike of the original virus, but they are ineffective against viruses that carry E484K or N501Y substitutions in the RBD. However, in multimeric form, these Nbs overcome the block and display remarkable neutralization potency. This reversal is likely the result of increased avidity for the trimeric spike, or possibly the simultaneous cross-linking of multiple spikes on the viral membrane.

The second group of Nbs (Nb12, Nb30) associates with a region that is highly conserved among coronaviridae (Wrapp et al., 2020) but remains inaccessible to most human antibodies. As this region lies outside the ACE2 binding motif, Nb-RBD contacts are unaffected by E484K or N501Y. Importantly, even though the conserved domain does not overlap with the ACE2 binding motif, our structural studies show that Nbs of this class sterically interfere with ACE2-RBD associations.

Overall, the Nbs described here solve the challenge posed by SARS-CoV-2 escape variants either through enhanced avidity as homotrimeric constructs or by binding conserved epitopes typically unavailable to human monoclonal antibodies. Combining the two Nb classes into heterotrimeric constructs might further improve their efficacy, as was recently shown with Nb permutations that prevented viral escape *in vitro* (Koenig et al., 2021). Finally, as is often the case with single-chain antibodies, the Nbs were thermostable and can be aerosolized with commercially available mesh nebulizers without losing neutralization activity (Figure S6B-D). Based on these features we propose that our leading Nbs may provide valuable tools for passive immunotherapy or pulmonary delivery against current and future SARS-CoV-2 variants of concern.

## Supporting information

This file contains all methods

## Figure Legends

**Figure S1:**
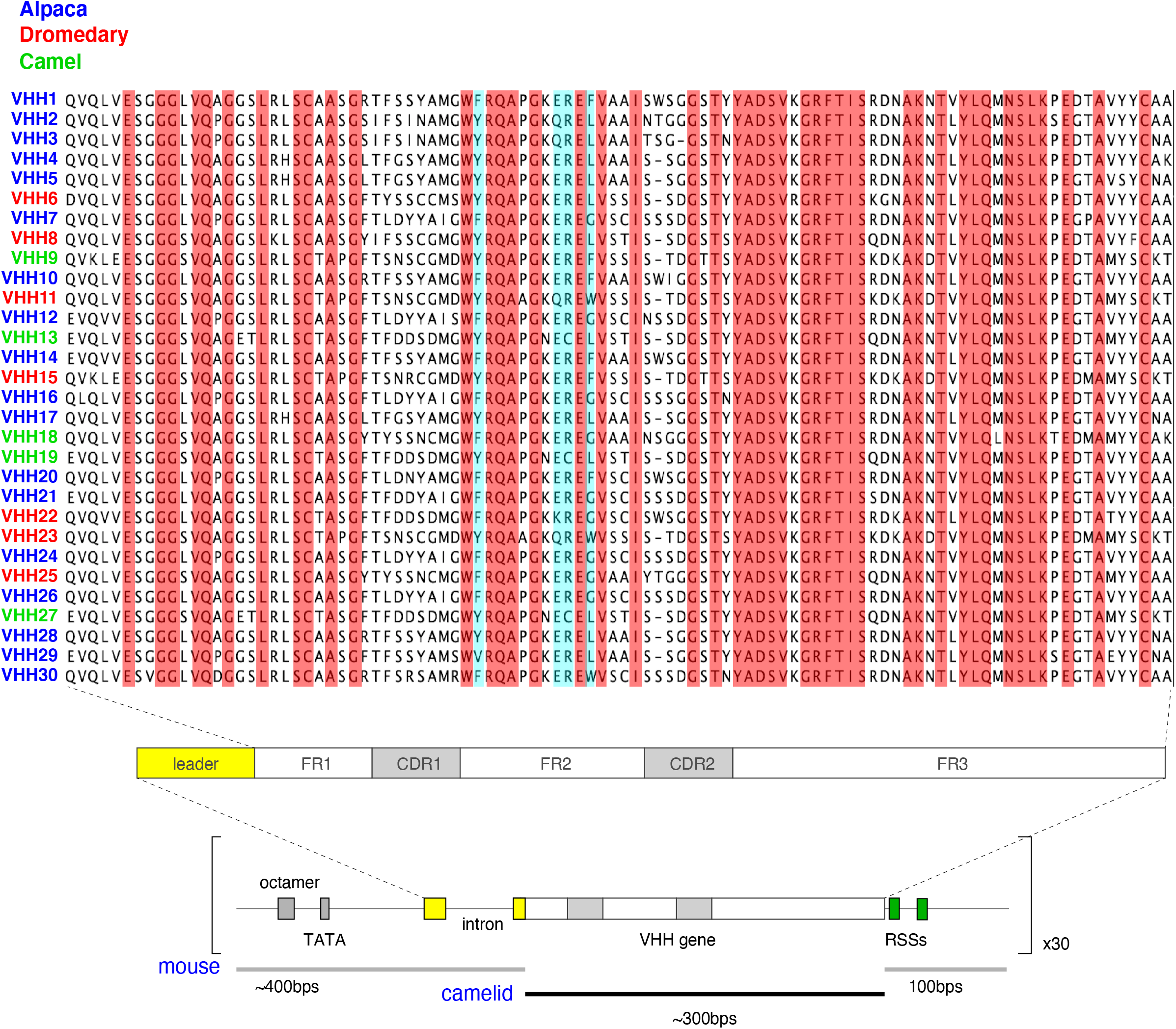
VHH genes used in the array and gene unit assembly. Alignment of the 30 VHH genes highlighting in red 100% AA conservation and in blue the 4 hydrophilic AAs in FR2 [in VH proteins, those 4 AAs are hydrophobic and mediate the interaction with light chains]. Schematics below show the configuration of VHH gene units, composed of a mouse VH promoter (250 bps containing the octamer and TATA box); mouse leader exons (∼150 bp) codifying for the signal peptide cleaved off during heavy chain processing in the ER; the camelid VHH ORF (∼300 bps); mouse downstream sequences (100 bps) containing the recombination signal sequences (RSSs).

**Figure S2:**
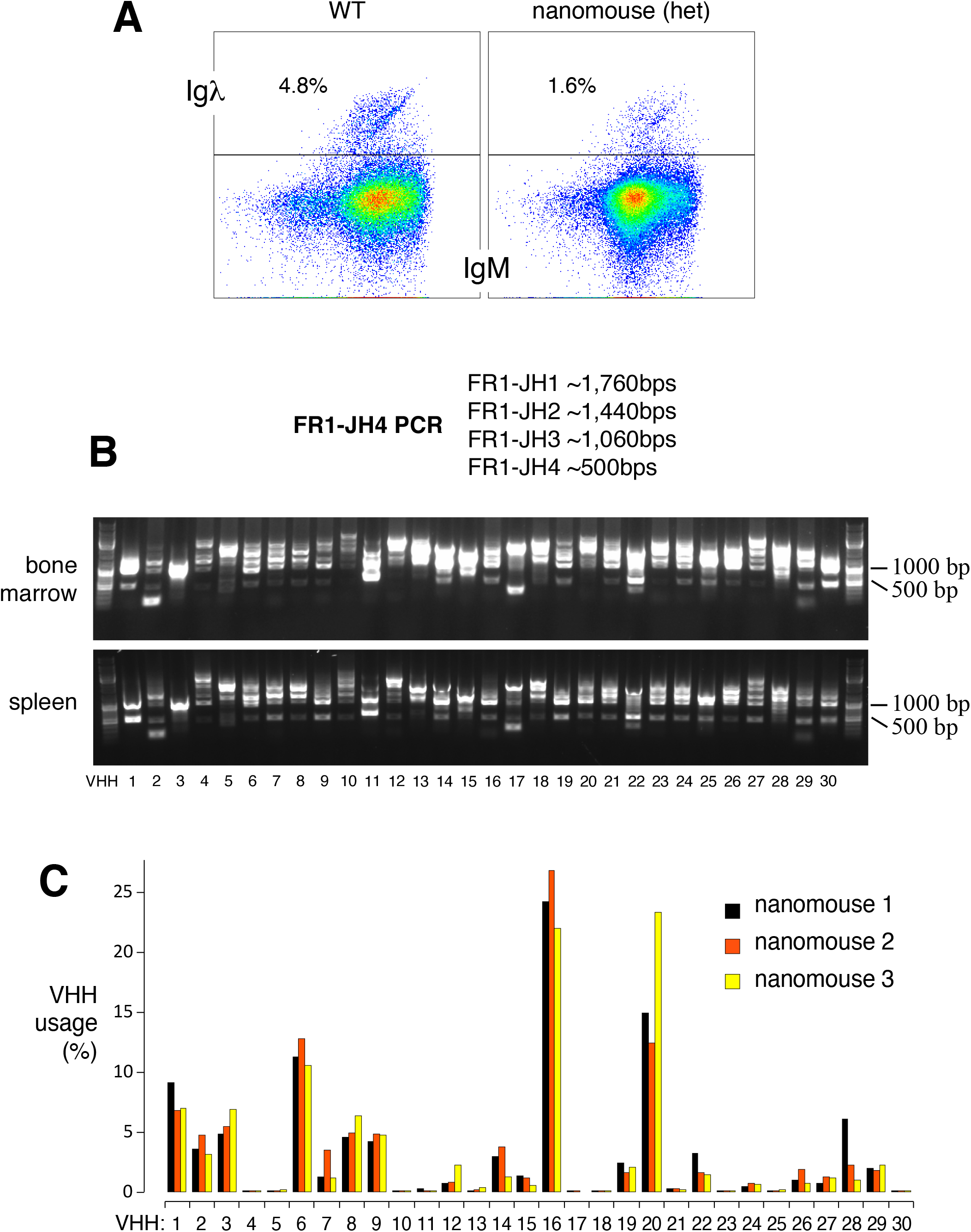
Igλ expression and recombination frequency of VHH genes. (**A**) Flow cytometry analysis of Igλ expression in B220^+^IgM^+^ splenic B cells from wild type and heterozygous nanomice. (**B**) VHH-DJ recombination was monitored by genomic PCR in bone marrow and spleen samples using a framework 1 (FR1) VHH-specific primer and a second primer downstream of JH4. The expected PCR products for each recombination event between a given VHH and JH1, JH2, JH3, or JH4 are provided. (**C**) Bar graph showing VHH percentage usage among splenic B cells in three nanomouse littermates.

**Figure S3:**
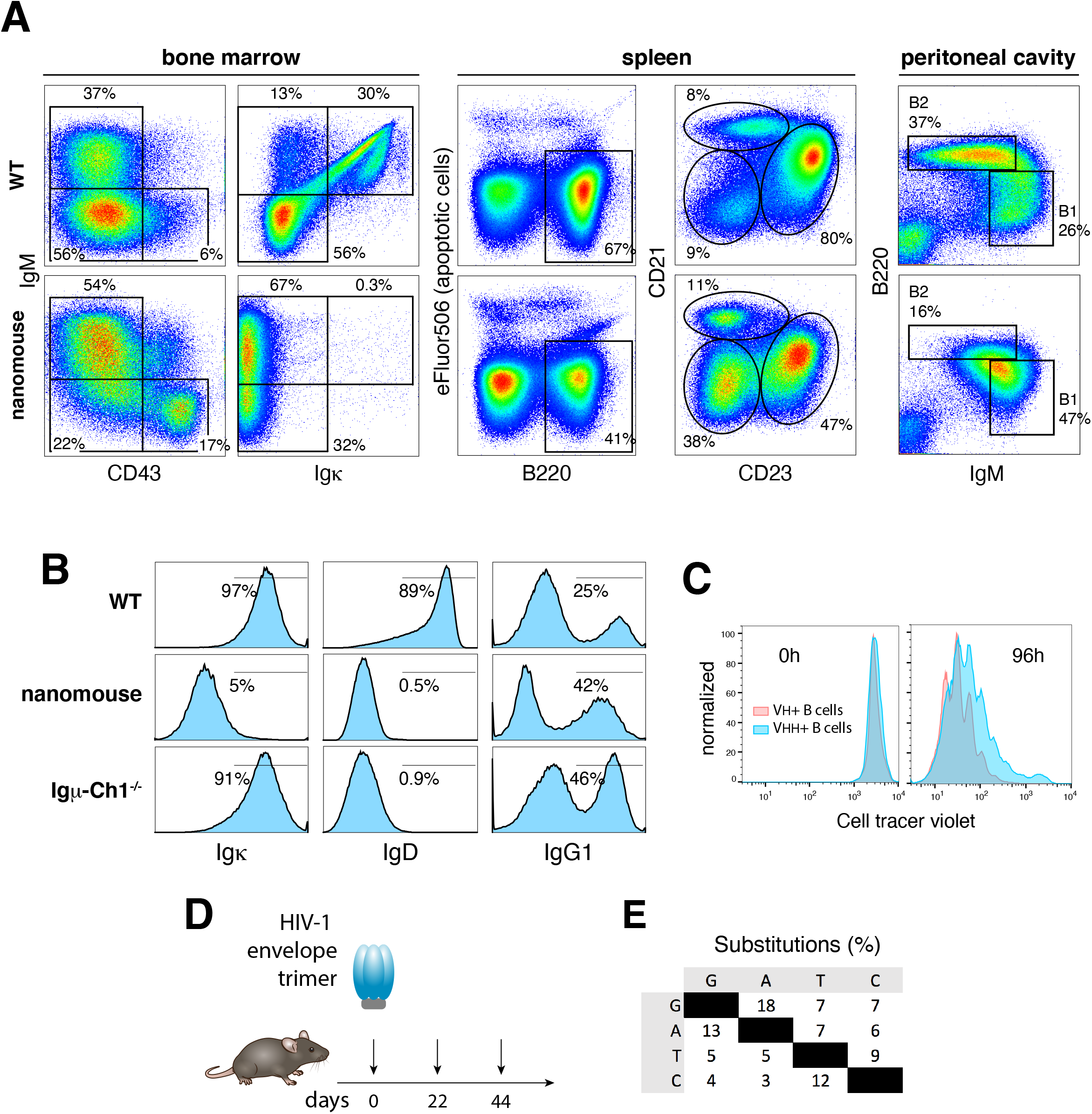
B cell development in nanomice. (**A**) Flow cytometry analysis of bone marrow (left two columns), spleen (third and fourth columns), and peritoneal cavity B cells (last column) in WT controls and nanomice. First column shows the percentage of B220-gated CD43^+^IgM^-^ proB and CD43^-^IgM^+^ immature B cells. Second column shows percentage of Igκ within the B220-gated IgM^+^ population. Third column denotes the total number of B220^+^ B cells in the spleen. Y axis shows viability staining with eFluor506 (eBiosciences). The fourth column shows the percentage of B220-gated CD23^low^CD21^low^ immature, CD23^high^CD21^low^ follicular, and CD23^low^CD21^high^ marginal zone splenic cells in the two strains. The last column shows the percentage of B1 (IgM^high^B220^low^) and B2 (IgM^low^B220^high^) cells in the peritoneal cavity. (**B**) Histograms depicting the percentage of Igκ (left), IgD (middle), and IgG1 (right row) in WT, nanomice, and Igµ-CH1^-/-^ mice. The latter measured in LPS+IL-4+αCD180 *ex-vivo* cultures. Population gates are represented with a line and the percentage of total cells is provided. (**C**) Proliferation assay of nanomouse and control B cells cultured for 96h with LPS+IL-4+αCD180. (**D**) Immunization regimen. Nanomice were immunized with 50 µg HIV-1 envelop trimer at the indicated dates. (**E**) Percent nucleotide substitutions (adjusted for base composition) observed in Nbs isolated from immunized animals. Phage library was selected for binding to HIV-1 envelop trimer.

**Figure S4:**
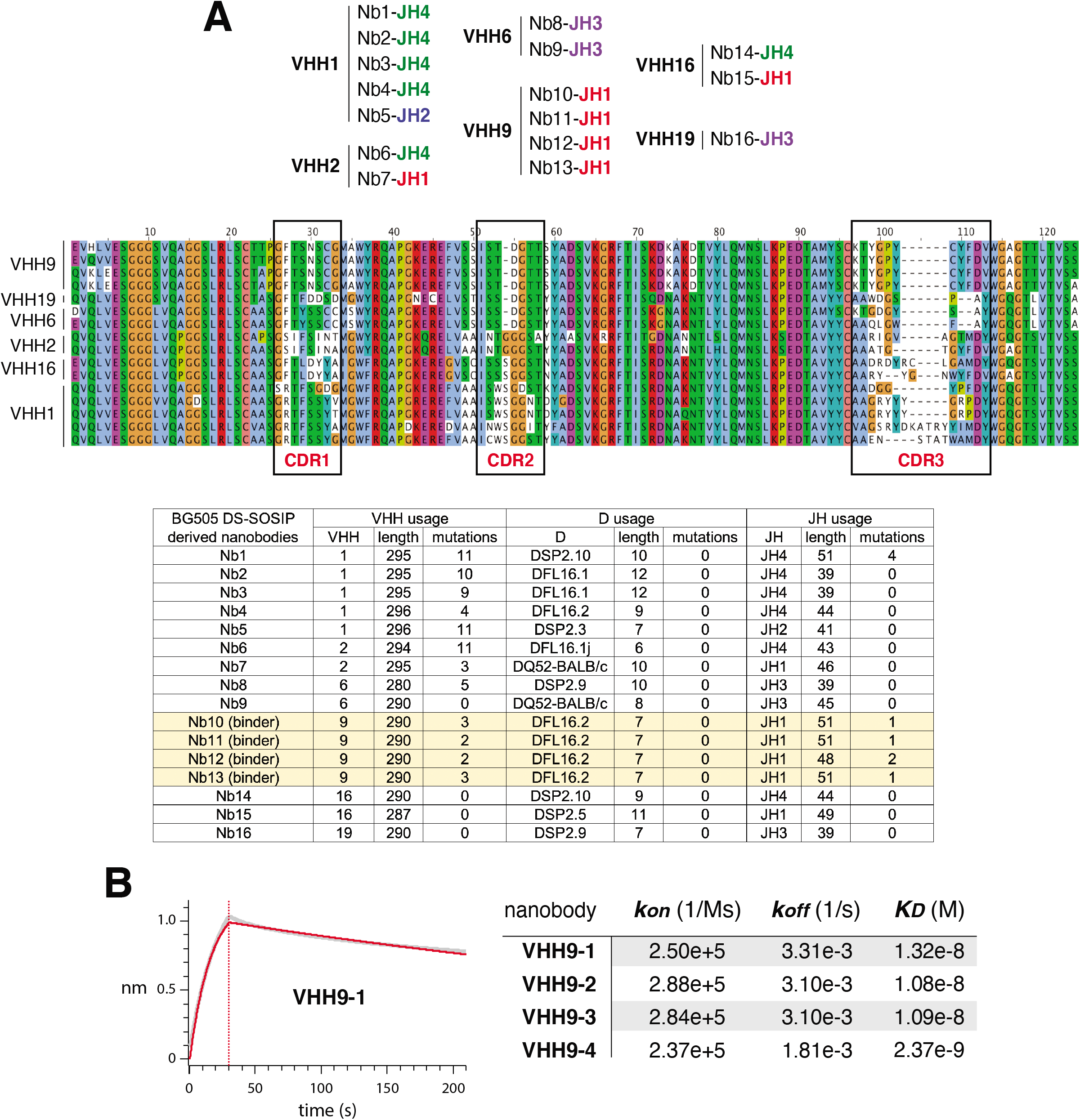
Nanomouse immune response to HIV-1 envelop trimer. (**A**) Upper table shows Nb and JH usage for the 16 VHH clones isolated from immunized nanomice. Middle graph shows protein alignment for VHHs isolated from HIV-1 trimer immunized nanomice. CDRs are boxed. Lower table shows hypermutation profiles for selected Nbs’ VHH, D, and J domains. (**B**) Left, BLI analysis of BG505 DS-SOSIP binding to immobilized VHH9-1. Red trace represents the raw data, and the kinetic fit is shown in grey underneath. Right, table showing the kinetic constants for association (*kon*), dissociation (*koff*) and equilibrium (*KD*) for all four VHH9 Nb variants.

**Figure S5:**
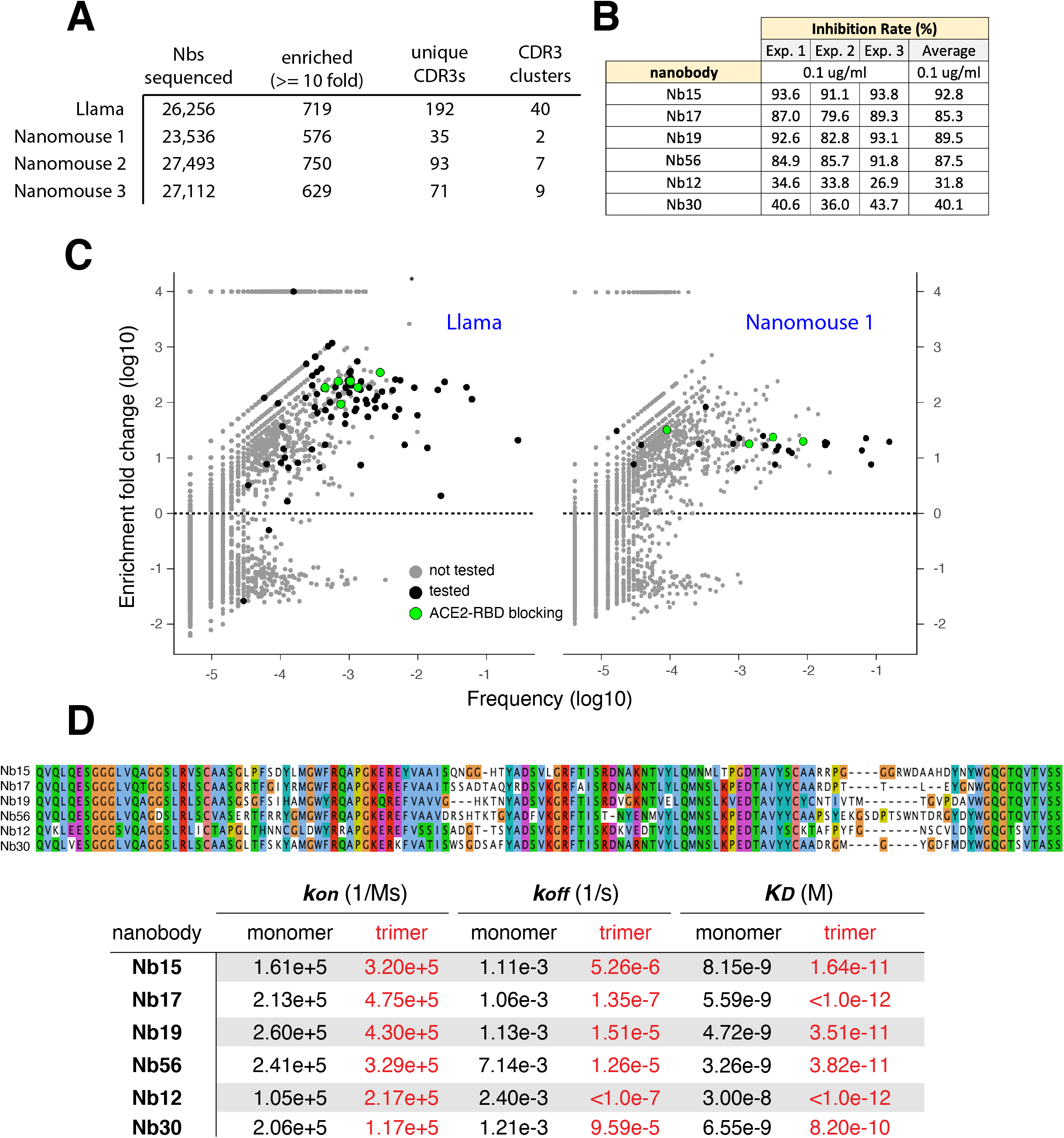
Isolation of anti-SARS-CoV-2 RBD Nbs. (**A**) Table indicating i) the total number of unique Nb genes identified from llama and three nanomice phage display libraries following selection for RBD binding, ii) the number of Nbs enriched at least 10-fold post-selection, iii) the number of Nbs with a unique CRD3, and iv) the different clusters of Nbs that share similar CDR3s (with no more than 2 AA differences). (**B**) Table showing *in vitro* neutralization results for the 6 leading Nbs using GenScript’s sVNT kit. (**C**) Dot plot depicting the extent of enrichment (y-axis) and frequency (x-axis) of unique Nbs post RBD selection of llama (left) or nanomouse 1 (right) libraries. Green circles represent Nbs that block ACE2-RBD interactions *in vitro*, black circles are Nbs that do not efficiently block ACE2-RBD interactions, and grey dots represent untested Nbs. (**D**) Upper graph shows protein alignment of the 6 Nbs isolated from llama and nanomice immunized with SARS-CoV-2 spike and RBD. Lower table shows equilibrium (*KD*), association (*Kon*) and dissociation (*Koff*) constants obtained for each Nb as a monomer (black) or trimer (red) form.

**Figure S6:**
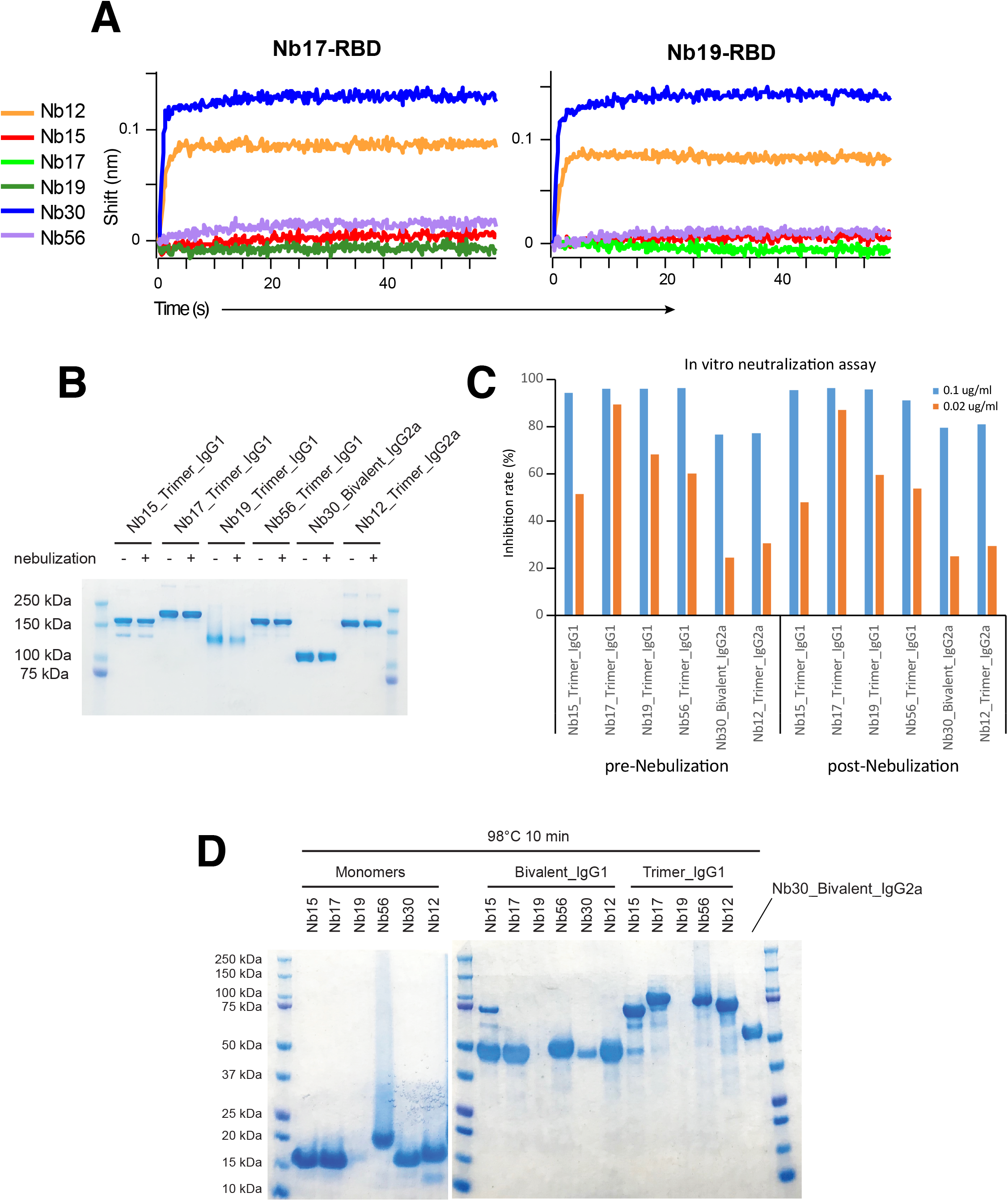
Nb competition for RBD binding and stability upon heat treatment and aerosolization. (**A**) Additional panels related to Figure 3c-f. Immunocomplexes used were Nb17-RBD (left) and Nb19-RBD (right). (**B**) Coomassie staining showing Nb integrity following nebulization. With the exception of Nb30 (bivalent) all Nbs were fused to Fcs as trimers. (**C**) Bar graph showing *in vitro* neutralization (percentage) of RBD-ACE2 interactions by the different Nb trimers (bivalent for Nb30) before and after nebulization at two different concentrations (0.1 µg/ml (blue) and 0.02 µg/ml). (**D**) Coomassie staining showing integrity of Nb monomers (left) or multimers (right) following heat treatment (98°C for 10 min).

**Figure S7:**
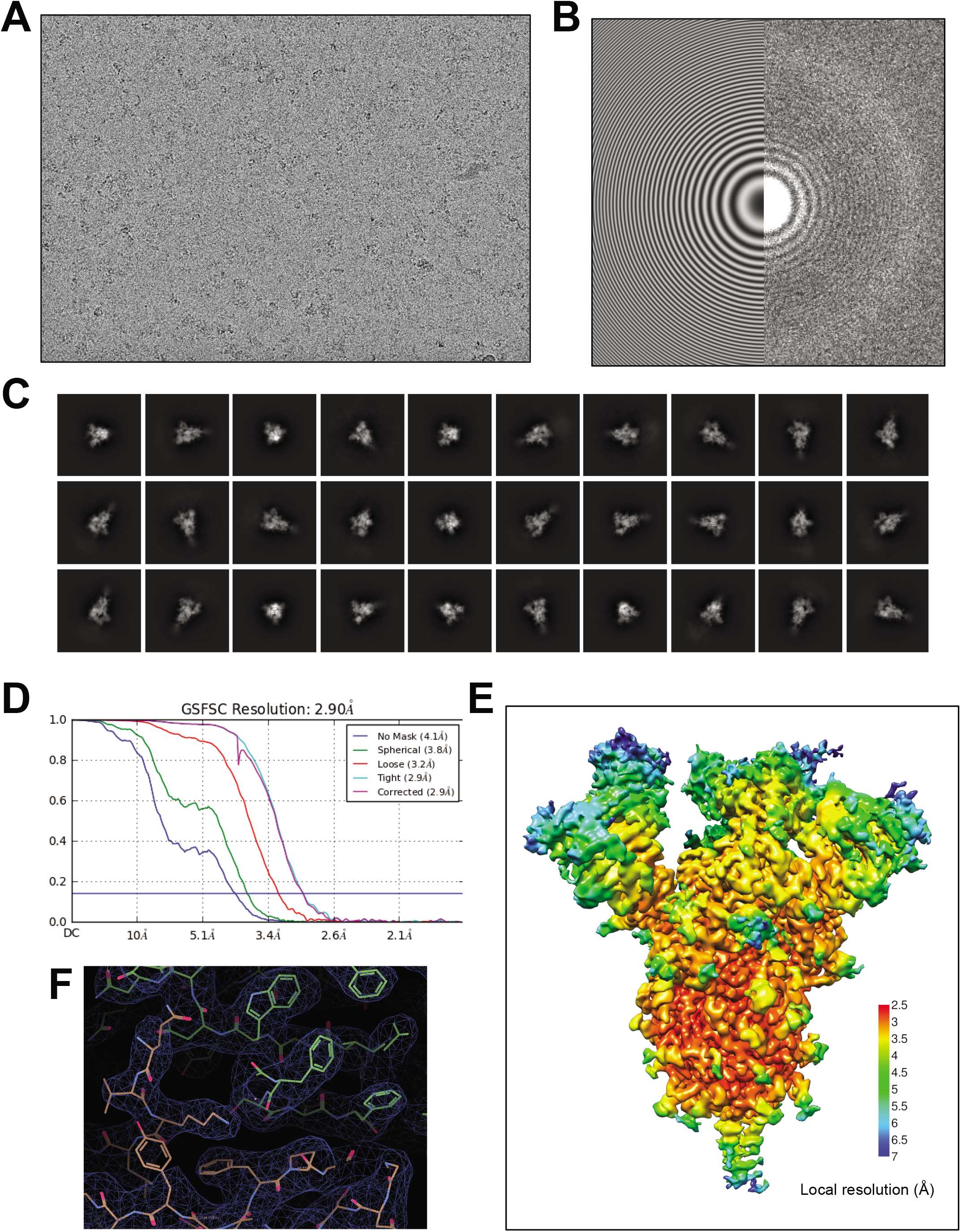
CryoEM data processing and validation for Nb12-spike complex. (**A**) A representative cryoEM micrograph showing Nb12-spike complex embedded in vitreous ice. A CTF fit of the micrograph. (**C**) Representative 2D average classes. (**D**) Overall Resolution estimation (FSC, 0.143). (**E**) Local resolution estimation of the cryoEM map. (**F**) CryoEM density and models for an interface region between RBD and Nb12 after local refinement.

**Figure S8:**
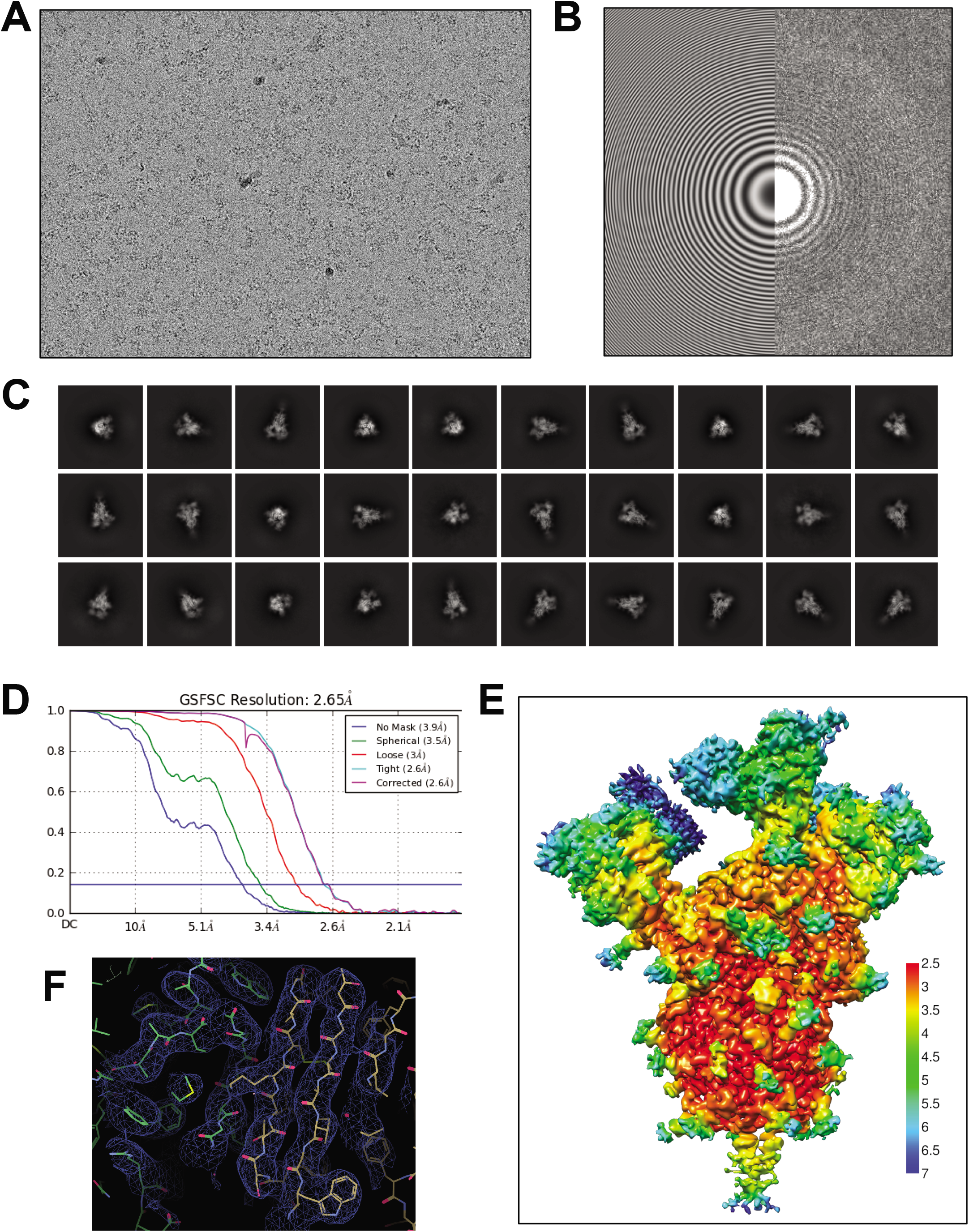
CryoEM data processing and validation for Nb30-spike complex. (**A**) A representative cryoEM micrograph showing Nb30-spike complex embedded in vitreous ice. (**B**) A CTF fit of the micrograph. (**C**) Representative 2D average classes. (**D**) Overall Resolution estimation (FSC, 0.143). (**E**) Local resolution estimation of the cryoEM map. (**F**) CryoEM density and models for an interface region between RBD and Nb30 after local refinement.

**Figure S9:**
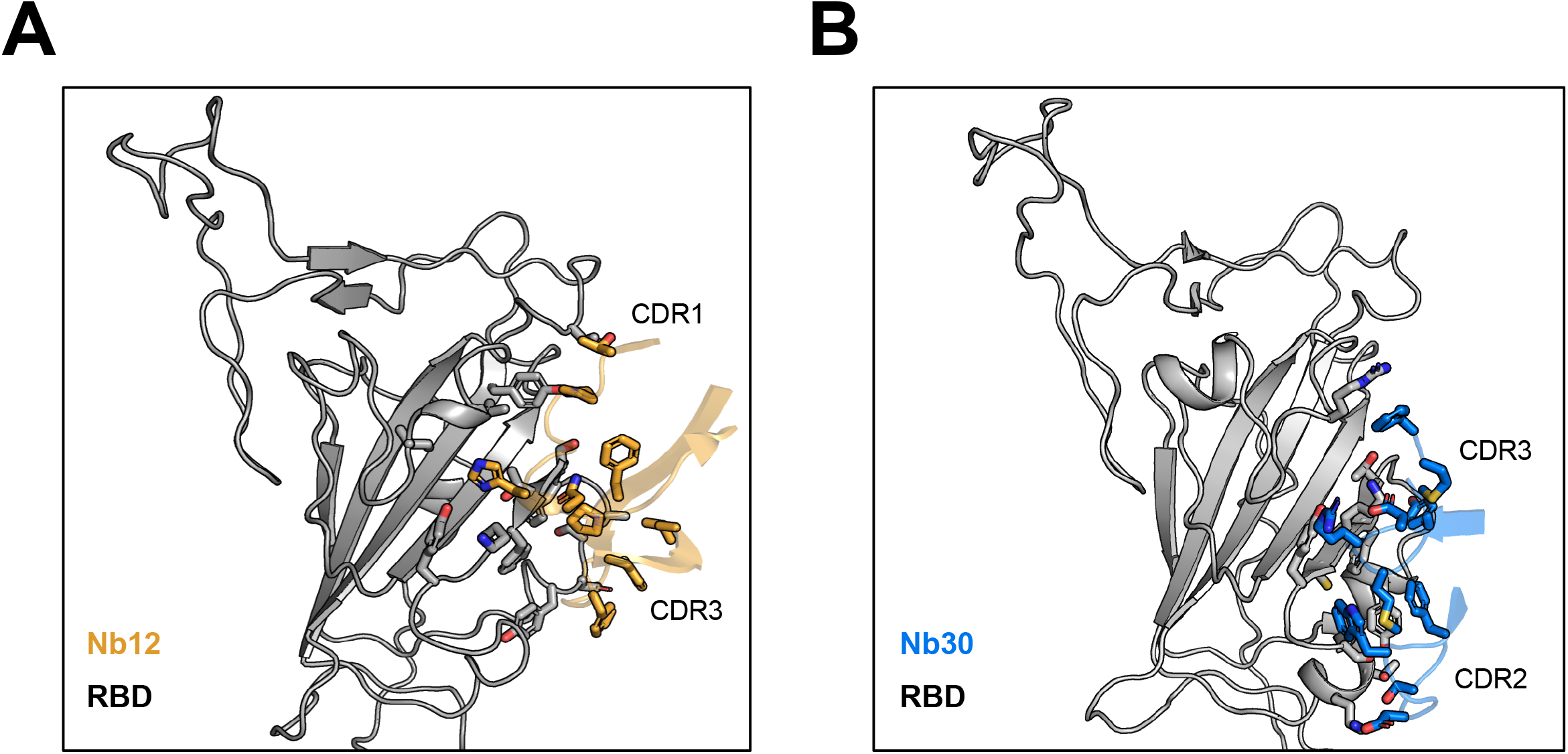
Structural analysis of nanomice and llama nanobody interaction to the SARS-CoV2 spike. (**A**) Interface between Nb12 and RBD. (**B**) Interface between Nb30 and RBD.

**Figure S10:**
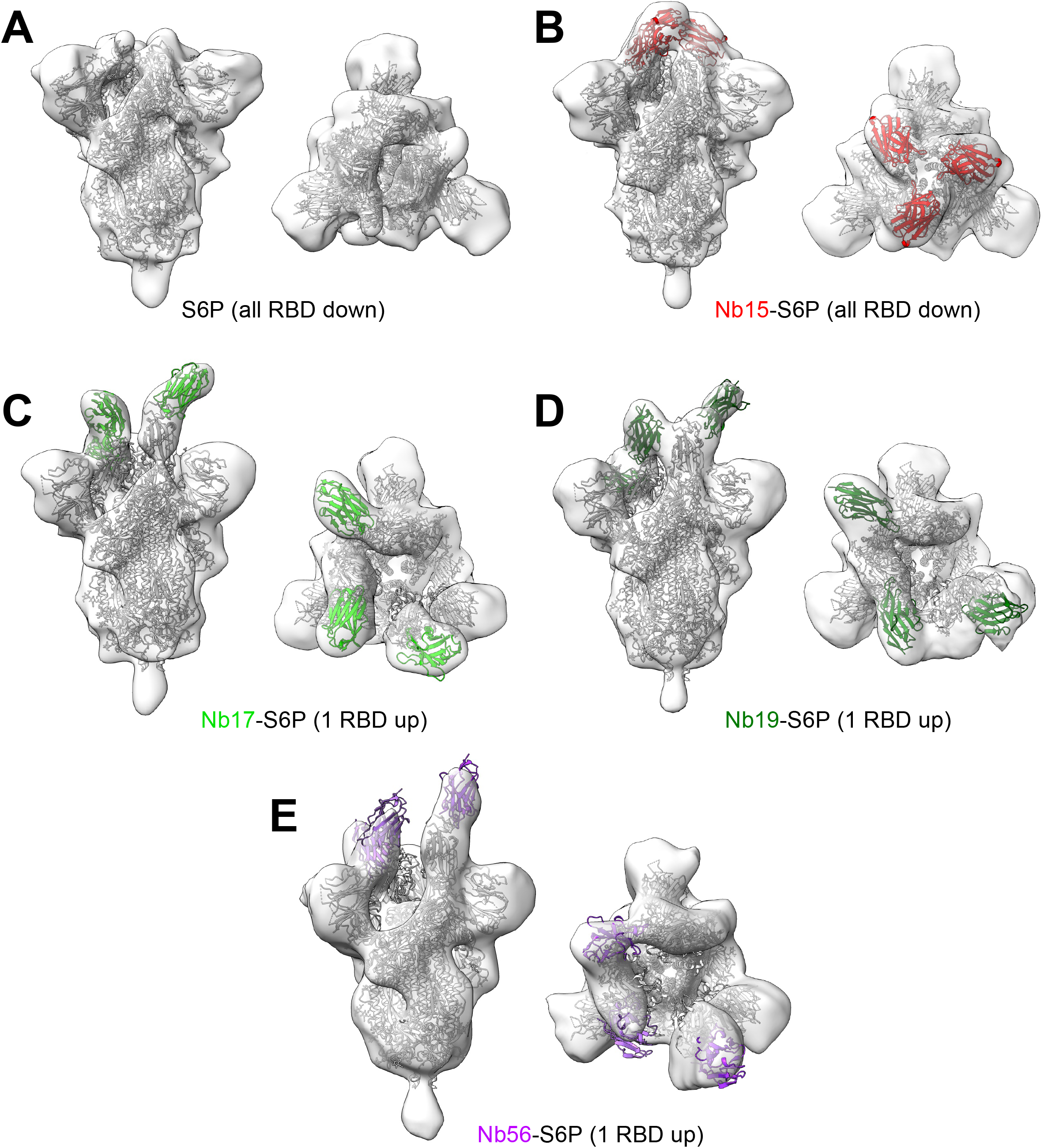
3D reconstruction of llama Nb-spike complex from negative stain EM. (**A**) SARS-CoV2 spike (Hexpro) structure in two perpendicular views. (**B**) Spike-Nb15 (red) complex structure in two perpendicular views. (**C**) Spike-Nb17 (light green) complex structure in two perpendicular views. (**D**) Spike-Nb19 (dark green) complex structure in two perpendicular views. (**E**) Spike-Nb56 (purple) complex structure in two perpendicular views.

