## Supplementary material for "Multimeric nanobodies from camelid engineered mice and llamas potently neutralize SARS-CoV-2 variants": This file contains all methods

### Data reporting

No statistical methods were used to predetermine sample size. The experiments were not randomized and the investigators were not blinded to allocation during experiments and outcome assessment.

### Construction of exchange-cassette and VHH minigene

To replace the entire mouse VH locus (mm10, chr12:113,567,224-116,010,427) we assembled a targeting vector (pLH28-exchange-cassette) and a VHH minigene. The targeting vector was built by inserting a selection cassette composed of pEF1a-Puro-TK-2A-EGFP in between wtLoxP and LoxP257 sites within the pLH28 exchange vector[1]. As homology arms, 1Kb and 0.8Kb fragments flanking the VH deletion domain were cloned 5' and 3' of the LoxP sites. To build the VHH minigene, VHH genes (18 from alpaca, 7 from dromedary and 5 from camel) were selected based on published sequences[2-4]. 30 mouse VH promoters (250 bps) were next chosen based on their activity (GRO-Seq) in resting and activated mouse B cells. VHHs were codon optimized and complemented with mouse leading exons, introns, and recombination signal sequences (RSSs). The 30 units were pieced together by Gibson assembly (NEB) into the pBeloBAC11 vector.

### ES cell targeting

E14 were cultured in Glasgow's MEM (Thermo Fisher Scientific, 11710035) supplemented with 10% FBS (ATCC, SCRR-03-2020), Glutamax, Sodium Pyruvate, NEAA, Pen/Str and β-mercaptoethanol (Thermo Fisher Scientific, 35050061, 11360070, 11140050, 15140122, 21985023, respectively) at 37 °C and 5% $CO_2$. mLIF (GeminiBio, 400-495, 10,000x), MEKi (Stemgent, 0400602, 10,000x) and GSKi (Stemgent, 0400402, 3,333x) were added to medium before use. Dishes and plates were coated with 0.1% glycine for 15 minutes at room temperature before use. To delete the CH1 exon of Cμ, sgRNAs targeting the flanking introns were cloned into CRISPR-Cas9 plasmid pX458 (Addgene, 48138). A 100nt long single-stranded oligonucleotide (ODN) donor (100μM, 3 μl) was co-transfected with the two Cas9-sgRNA plasmids (2 μg each) into ESCs (2 million cells) by Amaxa nucleofection kit (Lonza, VPH-1001, program A030). After 24 hours culture, GFP-high ESCs were FACS sorted and cultured in 10-cm dishes at the concentration of 2,000 cells per dish. 7 days later, colonies were transferred into 96-well plates and cultured for an additional 3 days. Genomic DNA was then extracted and genotyped for Cμ exon deletion. Clones with homozygous deletion were selected to delete the Cγ1 exon with the same strategy. To delete the entire VH locus, sgRNAs targeting sequences upstream of Ighv1-86 (first Ighv) and downstream of Ighv5-1 (last Ighv) respectively were cloned into pX458. Selected ESCs (2 million cells) were co-transfected with the two Cas9-sgRNA plasmids (1.5 μg each) and the pLH28-exchange-cassette plasmid (1.5 μg) and then cultured in 10-cm dishes. 24 hours later, cells were selected with puromycin (0.8 μg/ml) for 10 days and individual colonies were picked for expansion and genotyping by long-range PCR. Positive clones (2 million cells) were co-transfected with VHH minigene vector (3 μg) and a Cre-expressing plasmid (1 μg) and cultured in 10-cm dish for 3 days. Cells were then selected with ganciclovir (2 μg/ml) for 7 days before individual colonies were picked for expansion and genotyping. sgRNAs and ODN primers are listed in Extended Data Table 4.

### Generation of nanomice

Two modified ESCs clones with normal karyotyping were injected into C57BL/6 blastocysts, which were then transferred to the uteri of pseudopregnant C56BL/6 recipients. High percentage chimeras were mated to C57BL/6 mice and offspring were genotyped for VHH minigene knock-in and Cμ and Cγ1 exon deletion. One out of three chimeras produced germline transmitted F1 offspring. Three F1 male mice were backcrossed with C57BL/6 mice. F2 heterozygous mice were inbred to produce mice homozygous for all three loci modifications. Two F1 offspring from the same chimera were $Ig\mu Ch1^{-/-}$ but WT for VH and CH1 exon of Cγ1. These mice were used as controls for Extended Data Fig. 3b.

**Fluorescence activated cell sorting (FACS) analysis**

For B cell activation, B cells were cultured in RPMI 1640 supplemented with 10% FBS, HEPES, Sodium Pyruvate, NEAA, Pen/Str and β-mercaptoethanol at 37 °C and 5% $CO_2$ in the presence of lipopolysaccharide (LPS), interleukin-4 (IL-4) and αCD180 antibodies for 72 hours. For proliferation assay, cells were stained with CellTracer Violet (Thermo Fisher Scientific, C34557) at room temperature for 20 minutes before culturing for 96 hours. For all FACS staining, cells were incubated in FACS buffer (PBS, 2% FBS) at 4 °C for 20 minutes. Antibodies used for staining include: anti-B220-PerCP-Cy5.5 (eBioscience, 45-045-82), anti-B220-APC (Invitrogen, 17-0452-83), anti-IgM-APC (eBioscience, 17-5790-82), anti-Igκ-PE (BD Pharmingen, 559940), anti-Igκ-FITC(BD Pharmingen, 550003), anti-Igλ-FITC (BD Pharmingen, 553434), anti-IgG1-PE (BD Pharmingen, 550083), anti-IgG1-APC (BD Pharmingen, 550874), anti-IgD-FITC (BD Pharmingen, 553439), anti-CD95-PE (BD Pharmingen, 554258), anti-CD43-PE (BD Pharmingen, 553271), anti-CD23-PE (BD Pharmingen, 553139), anti-CD21-FITC (Biolegend, 123408), Viability Dye eFluor506 (Invitrogen, 1923275). Data were acquired using BD FACSCanto and FACSDiva software and analyzed with FlowJo software.

**Analysis of VHH(D)J recombination**

Genomic DNA from bone marrow or splenic samples was extracted with the DNeasy Blood & Tissue kit (Qiagen, 69506). VHH(D)J joints were PCR amplified from 100ng of DNA with a framework primer unique for each of the 30 VHHs, and a common downstream JH4 primer. PCR products were loaded onto 1% agarose gel to resolved them by size. Primers are listed in Extended Data Table 4.

**VHH(D)J recombinants phagemid library construction**

VHH(D)J phagemid libraries from unimmunized mice were constructed by first extracting RNA from nanomouse splenic samples with Trizol Reagent (Thermo Fisher Scientific, 15596026) and reverse transcribed to cDNA with SuperScript™ III First-Strand Synthesis SuperMix (Thermo Fisher Scientific, 18080400) according to manufacturer's instructions with some modifications.10 μg of total RNA was denatured and annealed with gene specific primer corresponding to CH2 of Ighμ gene. After elongation at 50 °C for 50 minutes, template switching oligonucleotide (TSO, 3'-propyl modified) linker was added to the 3' end of the first strand cDNA with 90 minutes incubation at 42 °C. The reaction was inactivated at 85 °C for 5 minutes and 2 μl of cDNA was used as template for VHH(D)J amplification by two-step PCR with CloneAmp HiFi PCR Premix (Takara, 639298). For the first-step PCR, unmodified TSO and Ighμ CH2-specific oligonucleotides were used. 30 ng of the first-step PCR product was then amplified with a primer mix of 30 forward primers corresponding to framework (FR1) of 30 VHH genes and 4 reverse primers corresponding to JH1~JH4. pMES4 phagemid (Addgene, 98223) was amplified with primers to introduce SfiI sites on both ends. VHH(D)J and pMES4 fragments were then digested with SfiI (NEB, R0123L) and ligated (100 and 200 ng respectively) with T4 ligase (NEB, M0202L) at 16 °C overnight. Ligation product was purified with DNA Clean & Concentrator (Zymo Research, D4014) and eluted into 12 μl of water. 3 μl of DNA was electroporated into 60 μl of TG1 cells (Lucigen, 60502-2) in 1.0 mm cuvette (HARVARD Apparatus, 450134) with BTX electroporation system ECM 630 at the setting of 25 μF, 200 Ohms, 1600 Volts. After 1 hour of recovery in 37 °C in shaker incubator, TG1 cells were plated on 5 of 10-cm LB Agar plates supplemented with 100 μg/ml carbenicillin (KD Medical, BPL-2400). Plates were placed in 37 °C bacteria incubator overnight and then bacteria scaped off plates and phagemid library DNA extracted with Zymo Plasmid Miniprep kit (Zymo Research, D4054). Primers are listed in Extended Data Table 4.

**Sanger sequencing for somatic hypermutation analysis**

VHH(D)J recombinants from splenic cells of two nanomice were PCR amplified as described in the phagemid library construction section, and then cloned directly into pCR-Blunt II-TOPO vector (Thermo Fisher Scientific, 450245) and transformed into Stabl3 competent E.coli (Thermo Fisher Scientific, C737303). 96 colonies were randomly picked for Sanger sequencing. TG1 cells from BG505 DS-SOSIP immunized nanomouse phagemid library were plated onto carbenicillin-containing plates and 50 colonies picked for Sanger sequencing.

**Immunizations**

All animal related procedures were performed by following our NIAMS ACUC protocol. To monitor the germinal center reaction, three nanomice and two C57BL/6 mice were immunized intraperitoneally with 50 μg of keyhole limpet hemocyanin (KLH) in the presence of complete Freund's adjuvant (CFA). A boost injection was performed in the footpads with 25 μg of KLH in the presence of incomplete Freund's adjuvant (IFA) on day 6. Spleen samples were harvested on day 12 for analysis.

To isolate nanobodies recognizing HIV-1 envelop trimer, one nanomouse was immunized intraperitoneally with 50 μg of BG505 DS-SOSIP in the presence of CFA on day 0, and boost immunized with 25 μg of BG505 DS-SOSIP in the presence of IFA or phosphate-buffered saline (PBS) on day 22 and 44, respectively. Bone marrow, spleen and blood were harvested on day 48.

To isolate neutralizing nanobodies against SARS-CoV-2, a llama (Capralogics, Inc.) was immunized subcutaneously with 1 mg of recombinant RBD protein in the presence of CFA at day 0, and boost immunized with 0.5 mg of RBD protein in the presence of IFA on day 14, 28, 42. Two more boost immunizations with 0.5 mg of recombinant Spike protein in the presence of IFA were performed on day 56 and 70. On day 80, 500 ml of whole blood were collected for library preparation.

To isolate SARS-CoV-2 neutralizing nanobodies from nanomice, two groups of mice (five mice in group 1 and six mice in group 2) were immunized with RBD and/or Spike protein and bleeds were collected after a 62-day immunization protocol. Mice were immunized intraperitoneally with 50 μg of RBD protein (group 1) or Spike protein (group 2) in the presence of CFA on day 0, and boost immunized intraperitoneally with 25 μg of RBD protein (group 1) or Spike protein (group 2) in the presence of IFA on day 14, 28 and 42. Mice were further immunized with 25 μg of Spike protein in PBS on day 56 and 59, intraperitoneally and intravenously, respectively. Bone marrow, spleen and blood were harvested on day 62. Best responders nanomouse 1 (group 1) and nanomouse 2 and 3 (group 2) were selected for phage library construction.

**Llama and nanomouse phage library construction**

The llama phage library was constructed as previously described[5] with some modifications. Briefly, 300 ml of whole blood was collected from llama and peripheral blood mononuclear cells (PBMC) were isolated using Ficoll-Paque plus (GE Healthcare, 17-1440-03). 50 μg of

extracted RNA was reverse transcribed to cDNA with random hexamers and 2.5 μl of cDNA was used for first round RT-PCR with gene-specific primers CALL001 and CALL002. The PCR reaction was repeated in 12 individual tubes with cDNA added into reactions separately. PCR fragments of about 700 bp long were gel purified and used as template (30 ng for each reaction, repeated in 12 individual tubes) for second round PCR with nested primers VHH-Back and VHH-For. PCR product from individual reactions were pooled and gel purified. Nanobody fragments and pMES4 phagemid were digested with PstI-HF and BstEII-HF restriction enzymes (NEB R3140L, R3162L) and ligated (1 μg and 2 μg respectively) with T4 ligase at 16°C overnight. Ligation product was column purified (into 12 μl of H2O) and electroporated into 360 μl of TG1 cells. After 1 hour of recovery in 37 °C in shaker incubator, cells were plated on 6 of 245mm x 245mm dish (Thermo Fisher Scientific, 431301) containing 2-YT Agar supplemented with 100 μg/ml carbenicillin and 2% (wt/vol) glucose. Plates were placed in 37°C bacteria incubator overnight and then bacteria scaped off plates and archived as glycerol stocks. Cells were infected with VCSM13 helper phage (Agilent Technologies, 200251) followed by precipitation of culture supernatant with 20% polyethylene glycol 8000 (Sigma, 89510) /2.5M sodium chloride on ice to purify the nanobody phage particles. Phage particles were resuspended in 1 ml PBS, 300 μl was used for screening immediately and the remaining phages were stored in -80°C in the presence of 10% glycerol.
Nanomouse nanobody phage libraries were constructed the same way as nanomouse VHH(D)J region phagemid library construction with some modifications. Briefly, total RNA was extracted from splenic cells, bone marrow cells and PBMC of immunized mice and processed separately until TG1 cell electroporation step. RNA from splenic cells, bone marrow and PBMC (50 μg, 50 μg and all respectively) were reversed transcribed to cDNA with Ighγ1 CH2-specific primer in separate tubes. 2 μl of cDNA was used as template for BCR variable domains amplification (12 reactions each), using unmodified TSO and Ighγ1 CH2-specific oligonucleotide as primers. Second PCR was repeated in 12 reactions using 30 ng of the first-step PCR product as template and 30 FR1 and 4 JH oligonucleotide mix as primers. PCR products were gel purified, digested with SfiI and ligated with pMES4 (200 ng and 400 ng respectively). Ligation products from splenic cells, bone marrow and PBMC samples were pooled and column purified (into 12 μl of water) and electroporated into 360 μl of TG1 cells and phage libraries prepared as described above. Primers are listed in Extended Data Table 4.

**Library construction for Illumina MiSeq deep sequencing**

Phagemid DNA extracted from TG1 cell libraries was used as starting material for constructing MiSeq libraries to measure VHH usage and nanobody diversity. Briefly, 1.2 μg of phagemid DNA was used as template and VHH(D)J inserts were amplified with primers recognizing the pMES4 backbone using CloneAmp HiFi PCR Premix (Takara, 639298) in a 50 μl reaction (9 cycles). To avoid MiSeq failure due to low complexity at initial cycles and to enable multiplex sequencing, 1-9 nt long staggers were introduced into forward primers. Without purification, 5 μl of the first PCR product was used as template for a second PCR (9 cycles) to add Illumina P5 and P7 primers on both ends. PCR product was then loaded onto a 2% agarose gel and the ~580 bp size band was purified with Zymoclean Gel DNA Recovery Kit (Zymo Research, D4002). DNA concentration was determined by Qubit 4 Fluorometer (Thermo Fisher Scientific, Q33238) and average DNA size was determined by TapeStation 4150 (Agilent). DNA was then adjusted to 2 nM in elution buffer containing 0.1% Tween-20. For unimmunized nanomice VHH(D)J library, DNA (2nM) from 3 mice were mix at 1:1:1 ratio and loaded for MiSeq run. For immunized llama and nanomice nanobody diversity analysis,

DNA (2nM) of pre-selection and post-selection library were mixed at 10:1 ratio first and then samples from individual animals were pooled at 1:1 ratio before loading for MiSeq sequencing. Primers are listed in Extended Data Table 4.

**Deep sequencing analysis**

For unimmunized nanomouse VHH usage analysis, pooled library from 3 mice were sequenced by MiSeq (pair end, 270 cycles x 2). Pair end reads were merged with NGmerge[6] with default settings. Nucleotides corresponding to pMES4 were trimmed using pTrimmer program[7], leaving clean VHH(D)J sequences in the merged reads. Reads with undetermined N nucleotides, low quality sequence or less than 300 nt in length were removed with the fastp program[8]. Fastq format sequences were converted to fasta format for further analysis. To calculate VHH usage, a BLAST database was built from a fasta format file vhh.exon.fa containing the exon sequence of all 30 VHH genes, using BLAST+. VHH(D)Js were then aligned to VHH genes using igblast program[9]. Alignment output file was simplified to retain only sequence ID and VHH(D)J recombination information.

For immunized llama and nanomouse nanobody diversity analysis, in total 8 libraries (pre- and post-selection) were sequenced by MiSeq (pair end, 300 cycles x 2). The 3' end low-quality sequences were trimmed using the Sickle program (v1.33, available at https://github.com/najoshi/sickle.). For different libraries, the minimum length of trimmed sequence was adjusted based on the length of staggers in the primers used for library construction. Paired sequences were merged by flash program (v1.2.11)[10] and translated. To extract nanobody sequences and to locate CDR3 region, we used ANARCI program[11] to annotate VHH genes with IMGT numbering. Protein sequences with greater than or equal to 100 amino acids in total and greater than or equal to 1 amino acid in CDR3 region were extracted for further analysis. Enrichment of individual sequences were calculated by comparing their frequencies in pre- and post-selection libraries. Sequences that were enriched more than 10 times and had greater than or equal to 5e-05 frequency were selected for CDR3 clustering using cd-hit program (v4.6.8)[12].

**Expression and purification of BG505 DS-SOSIP and SARS-CoV-2 proteins**

BG505 DS-SOSIP protein was expressed and purified as described[13]. The spike protein of SARS-CoV-2 and its receptor binding domain (RBD) were expressed and purified as described[14,15] with some modifications. Briefly, 1 mg of pCAGGS-Spike or pCAGGS-RBD plasmid was transfected into 1 liter of Expi293 cells (Thermo Fisher Scientific, A14528) with Turbo293 transfection reagent (Speed Biosystem, PXX1002). Supernatants from transfected cells were harvested on day 4 post-transfection by centrifugation of the culture at 12,000 g for 15 min. Supernatant was then filtered through 0.2 m aPES filter (Thermo Fisher Scientific, 5670020) and incubated with 10 ml of cOmplete His-tag purification resin (Roche, 50434600) for 1 hour at room temperature. Next, His-tag resin was collected through gravity flow columns (BioRad, 9704652), washed with 100 ml of washing buffer (15 mM imidazole, 50 mM TrisHCl, 300 mM NaCl) and eluted with 25 ml of elution buffer (300 mM imidazole, 50 mM TrisHCl, 300 mM NaCl). Eluate was concentrated in 10 kDa Amicon Centrifugal Units (EMD Millipore, UFC901024) and then dialyzed in PBS using Slide-A-Lyzer dialysis cassette (Thermo Fisher Scientific, 66381). Proteins were analyzed by NuPAGE gel (Thermo Fisher Scientific, NP0336BOX) and visualized by InstantBlue staining (Abcam, ab119211). Soluble spike trimers or monomeric RBD proteins were aliquoted, snap frozen by liquid nitrogen and stored at -80°C before used for animal immunization. RBD and Spike (HexaPro) proteins used for phage screening, Bio-layer Interferometry assay, Negative-Stain and Cryogenic Electron Microscopy were done as previously described[16,17].

**Phage screening for BG505 DS-SOSIP, RBD and Spike binding Nbs**
RBD, Spike and BG505 DS-SOSIP were coated by different methods onto MaxiSorp 96-well plate (Thermo Fisher Scientific, 439454) for phage screening. For RBD screening, two wells were coated with 50 μl of RBD protein (100 μg/ml in PBS) in 4°C overnight. Another well with 50 μl of PBS was used as an un-coated control. Wells were washed with PBS with 0.1% Tween-20 three times and blocked with 5% non-fat milk in PBS at room temperature for 1 hour. For Spike or BG505 DS-SOSIP screening, three wells were coated with 50 μl of Lectin (EMD Millipore, L8275, 100 μg/ml in PBS) in 4°C overnight. Wells were washed and blocked with 10% non-fat milk in room temperature for 1 hour. After three washes, 50 μl of 100 μg/ml BG505 DS-SOSIP or Spike were added to two wells, incubated at room temperature for 2 hours and washed. The third well contained PBS and served as an un-coated control. 300 μl of phage particles was mixed with 300 μl of 10% non-fat milk and rotated gently at room temperature for 1 hour. 150 μl of blocked phage particles were then added into each well and incubated in room temperature for 2 hours with gentle shaking. After 15 washes, phages were eluted with TrypLE Express Enzyme (Thermo Fisher Scientific, 12605010) by shaking plates at 700 rpm at room temperature for 30 minutes and used immediately for selection efficiency estimation (10 μl of phage eluate) and recovery infection (the remaining eluate) as described[5]. Anti-RBD libraries were selected with RBD protein once, and libraries constructed from BG505 DS-SOSIP or Spike immunized animals were selected with BG505 DS-SOSIP or Spike (HexaPro) proteins twice.

**ELISA selection of anti-BG505 DS-SOSIP and anti-RBD Nbs**
After one or two rounds of selection, recovered TG-1 cells were plated and colonies were picked to prepare periplasmic extracts containing crude nanobodies for ELSA. Briefly, individual colonies were picked and grown in 96 deep-well plates (Thermo Fisher Scientific, 278743) in 2YT medium supplemented with 100 μg/ml of Carbenicillin and 0.1% Glucose. IPTG (final 1mM) was added when OD600 reached 1 and protein expression was induced in 30°C for 16 hours. Periplasmic extracts were prepared by resuspending bacteria pellet in 200 μl of PBS and rapid freezed in liquid nitrogen. Frozen cells were thawed slowly at room temperature and centrifuged at 4100 g for 15 minutes. Maxisorp plates were coated with lectin (2 μg/ml) followed by BG505 DS-SOSIP (2 μg/ml) or with RBD (2 μg/ml). After blocking, 100 μl of Nb-containing supernatant was transferred to the plates and incubated for 2 hours at room temperature. Plates were washed and then incubated with horse radish peroxidase (HRP) conjugated goat anti-alpaca VHH domain specific antibody (Jackson ImmunoResearch, 128-035-232) for 1 hour at room temperature. Plates were washed and then developed by addition of 50 μl of tetramethylbenzidine (TMB) (Thermo Fisher Scientific, 34028) for 10 minutes, then the reaction was stopped by adding 50 μl of 1M H2SO4. Absorbance at 450 nm was measured with Synergy microplate reader (BioTek).

**Expression and purification of nanobodies**
Phagemids from lead candidates identified by ELISA were extracted from TG-1 cells and transformed into WK6 cells (ATCC, 47078). Cultures were grown in 30ml of 2YT medium (100 μg/ml of carbenicillin and 0.1% glucose) at 37°C and 220rpm until OD600 reached 1. Protein expression was induced by 1mM IPTG at 30°C for 16 hours and then cells pelleted at 4100 g for 15 minutes. The resulting pellets were resuspended in 1 ml of PBS plus 30 μl of 0.5 MU/ml polymyxin B (Sigma, P1004) and incubated at 37°C with shaking for 1 hour. Cell debris were pelleted at 12,000 g for 5 minutes and nanobodies in the supernatant were purified using Capturem His-tagged purification kit (Takara, 635710). For larger scale of

nanobody production (0.2 to 1 liter of culture), nanobodies in the supernatant were purified by cOmplete His-tag purification resin and dialyzed in PBS as described above. Proteins were filtered sterile by 0.22 μm PVDF membrane (EMD Millipore, UFC30GVNB) before used for downstream assay.

**Expression and purification of Fc conjugated nanobodies in Expi293 cells**
Monomeric or trimeric nanobody sequences were fused to the Fc region of human IgG1 with 6xHis tag at the C terminal end and cloned into pVRC8400 vector. In trimeric form, nanobody units were connected through (GGGGS)x3 flexible linkers. In some cases, llama IgG2a hinge region was used in lieu of human IgG1 hinge. The Fc-fusion constructs were expressed in Expi293 cells as described above at 33°C from day 2 to day 4. Antibodies in the supernatant were purified using either His-tag (Roche, 05893801001) or protein A (Thermo Fisher Scientific, A26457). When protein A resin was used, antibodies were eluted by IgG elution buffer (Thermo Fisher Scientific, 21009) and brought to neutral pH by adding 1/10 volume of Tris-HCl (1M, pH 8). Antibodies were concentrated, dialyzed and filtered.

**SARS-CoV-2 surrogate virus neutralization test (sVNT)**
RBD-ACE2 interaction blocking potential of nanobodies was tested using the SARS-CoV-2 surrogate virus neutralization test kit (Genscript, L00847) according to the manufacturer's instructions. Briefly, HRP-RBD was diluted and incubated with specified concentrations of nanobodies for 30 minutes at 37°C. Samples were then transferred onto ACE2-coated plates and incubated for 15 minutes at 37°C. Plates were washed, and the assay was developed using TMB reagent and quenched with stop solution. Absorbance at 450 nm was measured with a Synergy microplate reader (BioTek).

**Pseudotyped virus neutralization assay**
A panel of plasmids expressing RBD-mutant SAR-CoV-2 spike proteins in the context of pSARS-CoV-2-SD19 have been described previously[18,19]. The mutants E484K and KEN (K417N+E484K+N501Y) were constructed in the context of a pSARS-CoV-2-S$_{\Delta19}$ variant with a mutation in the furin cleavage site (R683G). The IC50 of these pseudotypes were compared to a wildtype SARS-CoV-2 spike sequence carrying R683G in the subsequent analyses, as appropriate. Generation of SARS-CoV-2 pseudotyped HIV-1 particles and pseudovirus neutralization assay was performed as previously described[20]. Briefly, 293T cells were transfected with pNL4-3DEnv-nanoluc and pSARS-CoV-2-SD19 and pseudotyped virus stocks were harvested 48 hours after transfection, filtered and stored at -80°C. Serially diluted nanobodies were incubated with the pseudotyped virus for 1 h at 37°C. The mixture was added to 293T$_{ACE2}$[21] (for analysis of WT neutralization activity, Figure 2) or HT1080Ace2 cl.14[22] cells (for analysis of spike mutant panel, Figure 3) and after 48 hours cells were washed with PBS and lysed with Luciferase Cell Culture Lysis 5x reagent (Promega). Nanoluc Luciferase activity in lysates was measured using the Nano-Glo Luciferase Assay System (Promega). The relative luminescence units were normalized to those derived from cells infected with SARS-CoV-2 pseudotyped virus in the absence of antibodies. The half-maximal inhibitory concentration for nanobodies (IC50) was determined using four-parameter nonlinear regression (GraphPad Prism).

**Nanobody stability studies**
Nanobody was nebulized with a portable mesh nebulizer (Philips, InnoSpire Go) producing 2-5 μm particles at a final concentration of 0.4 mg/ml. The resulting aerosol was collected by condensation into a 50 ml tube cooled on ice. Pre- and post-nebulization samples were analyzed by NuPAGE gel and visualized by InstantBlue staining. SARS-CoV-2 surrogate

virus neutralization test was also performed to compare the neutralization potency of pre- and post-nebulization samples. For thermostability test, nanobodies supplemented with loading buffer (Thermo Fisher Scientifc, NP0007) and β-mercaptoethanol were heated at 98°C for 10 minutes and then analyzed on NuPAGE gel and visualized by InstantBlue staining.

**Bio-layer Interferometry assay for nanobody affinity measurement**
Bio-Layer Interferometry (BLI) assay was performed using a fortéBio Octet Red384 instrument to determine the affinity of nanobodies to RBD. Briefly, biotinylated-RBD was immobilized onto streptavidin coated biosensors and then dip into nanobody solution for association for 30 seconds followed by dissociation for 2-3 minutes. Sensorgrams of the concentration series were corrected with corresponding blank curves and fitted globally with Octet evaluation software using a 1:1 Langmuir model of binding.

**Nanobody-RBD binding competition assay**
Nanobody-RBD binding competition assay was performed using a fortéBio Octet Red384 instrument. Biotinylated-RBD was first immobilized onto streptavidin coated biosensors and allowed association with one of the six nanobodies, then dip into one of the other five nanobodies. The competition of RBD binding can be determined by the absence of increasement in response unit.

**Negative-staining EM analysis for the structure of nanobody-spike complex**
Nanobody-spike complex samples were prepared by manually mixing two proteins in a 1:1 weight ratio, then diluted with a buffer containing 10 mM HEPES, pH 7.4, 150 mM NaCl, adsorbed to a freshly glow-discharged carbon-coated copper grid, washed with the above buffer, and stained with 0.75% uranyl formate. Images were collected at a magnification of 57,000 using EPU on a Thermo Fisher Talos F200C microscope equipped with a 4k x 4k CETA 16M camera and operated at 200 kV. The pixel size was 2.5 Å for the CETA camera. Particles picking, reference-free 2D classification, 3D reconstruction and refinement were performed using cryoSPARC.

**Cryo-EM data collection and processing**
Nanobody-spike complexes (Nb12-S6P and Nb30-S6P) were prepared by manual mixture of the two proteins in a 1:1 weight ratio, then diluted to a final concentration of 0.5 mg/ml. Sample (2.7 μl) was applied to a glow-discharged Quantifoil R 2/2 gold grids and vitrified using a Vitrobot Mark IV with a blot time of 3 s before the grid was plunged into liquid ethane. Data were acquired using the Leginon system installed on Titan Krios electron microscopes operating at 300kV and equipped with K3-BioQuantum direct detection device. The dose was fractionated over 40 raw frames and collected over a 2 s exposure time. Motion correction, CTF estimation, particle picking, 2D classifications, ab initio model generation, heterogeneous refinements, 3D variability analysis and homogeneous 3D refinements were carried out in cryoSPARC. Local refinement was performed to resolve the RBD-Nb interface by using a mask encompassing one copy of the RBD-Nb complex for refinement, after removing the rest density by particle subtraction.

**Cryo-EM model fitting**
For initial fits to the cryo-EM reconstructed maps, we used the coordinates of SARS-CoV2 spike from PDB ID 7JZL, and nanobody model predicted by ABodyBuilder server[23]. These initial models were docked into the cryo-EM maps using Chimera. The coordinates were then fit to the electron density more precisely through an iterative process of manual fitting using Coot and real space refinement within Phenix, Molprobity and EMRinger were used to check geometry and evaluate structures at each iteration step. Figures were generated in UCSF

ChimeraX and PyMOL (https://pymol.org). Map-fitting cross correlations were calculated using Fit-in-Map feature in UCSF Chimera. Overall and local resolution of cryo-EM maps was determined using cryoSPARC.

**Informatics analysis**

Sequence entropy are based on 9 strains with the following uniprot ID: SARS-Cov-2: P0DTD1, B.1.1.7, B.1.351; SARS-CoV: A7J8L4, Q202E5, P59594; and Bat SARS-like coronavirus: MG772933, Q0Q475, Q3I5J5. The entropy was calculated for each residue based on aligned sequences with the formula:

Entropy=$-\sum_{i=1}^{21} p(xi)\log(p(xi))$, where $x_i$ are standard amino acids, plus gap.

The buried surface area on the RBD were calculated for 51 human antibody/SARS-CoV-2 RBD complexes based using the Naccess program.


1 Han, L., Masani, S. & Yu, K. Overlapping activation-induced cytidine deaminase hotspot motifs in Ig class-switch recombination. *Proc Natl Acad Sci U S A* **108**, 11584-11589, doi:10.1073/pnas.1018726108 (2011).

2 Achour, I. *et al.* Tetrameric and homodimeric camelid IgGs originate from the same IgH locus. *J Immunol* **181**, 2001-2009, doi:10.4049/jimmunol.181.3.2001 (2008).

3 Bactrian Camels Genome, S. *et al.* Genome sequences of wild and domestic bactrian camels. *Nature communications* **3**, 1202, doi:10.1038/ncomms2192 (2012).

4 Wu, H. *et al.* Camelid genomes reveal evolution and adaptation to desert environments. *Nature communications* **5**, 5188, doi:10.1038/ncomms6188 (2014).

5 Pardon, E. *et al.* A general protocol for the generation of Nanobodies for structural biology. *Nat Protoc* **9**, 674-693, doi:10.1038/nprot.2014.039 (2014).

6 Gaspar, J. M. NGmerge: merging paired-end reads via novel empirically-derived models of sequencing errors. *BMC Bioinformatics* **19**, 536, doi:10.1186/s12859-018-2579-2 (2018).

7 Zhang, X. *et al.* pTrimmer: An efficient tool to trim primers of multiplex deep sequencing data. *BMC Bioinformatics* **20**, 236, doi:10.1186/s12859-019-2854-x (2019).

8 Chen, S., Zhou, Y., Chen, Y. & Gu, J. fastp: an ultra-fast all-in-one FASTQ preprocessor. *Bioinformatics* **34**, i884-i890, doi:10.1093/bioinformatics/bty560 (2018).

9 Ye, J., Ma, N., Madden, T. L. & Ostell, J. M. IgBLAST: an immunoglobulin variable domain sequence analysis tool. *Nucleic Acids Res* **41**, W34-40, doi:10.1093/nar/gkt382 (2013).

10 Magoc, T. & Salzberg, S. L. FLASH: fast length adjustment of short reads to improve genome assemblies. *Bioinformatics* **27**, 2957-2963, doi:10.1093/bioinformatics/btr507 (2011).

11 Dunbar, J. & Deane, C. M. ANARCI: antigen receptor numbering and receptor classification. *Bioinformatics* **32**, 298-300, doi:10.1093/bioinformatics/btv552 (2016).

12 Li, W. & Godzik, A. Cd-hit: a fast program for clustering and comparing large sets of protein or nucleotide sequences. *Bioinformatics* **22**, 1658-1659, doi:10.1093/bioinformatics/btl158 (2006).

13 Chuang, G. Y. *et al.* Structure-Based Design of a Soluble Prefusion-Closed HIV-1 Env Trimer with Reduced CD4 Affinity and Improved Immunogenicity. *J Virol* **91**, doi:10.1128/JVI.02268-16 (2017).
14 Amanat, F. *et al.* A serological assay to detect SARS-CoV-2 seroconversion in humans. *Nat Med* **26**, 1033-1036, doi:10.1038/s41591-020-0913-5 (2020).
15 Stadlbauer, D. *et al.* SARS-CoV-2 Seroconversion in Humans: A Detailed Protocol for a Serological Assay, Antigen Production, and Test Setup. *Curr Protoc Microbiol* **57**, e100, doi:10.1002/cpmc.100 (2020).
16 Zhou, T. *et al.* Structure-Based Design with Tag-Based Purification and In-Process Biotinylation Enable Streamlined Development of SARS-CoV-2 Spike Molecular Probes. *Cell Rep* **33**, 108322, doi:10.1016/j.celrep.2020.108322 (2020).
17 Hsieh, C. L. *et al.* Structure-based design of prefusion-stabilized SARS-CoV-2 spikes. *Science* **369**, 1501-1505, doi:10.1126/science.abd0826 (2020).
18 Weisblum, Y. *et al.* Escape from neutralizing antibodies by SARS-CoV-2 spike protein variants. *eLife* **9**, doi:10.7554/eLife.61312 (2020).
19 Wang, Z. *et al.* mRNA vaccine-elicited antibodies to SARS-CoV-2 and circulating variants. *Nature*, doi:10.1038/s41586-021-03324-6 (2021).
20 Robbiani, D. F. *et al.* Convergent antibody responses to SARS-CoV-2 in convalescent individuals. *Nature* **584**, 437-442, doi:10.1038/s41586-020-2456-9 (2020).
21 Robbiani, D. F. & Nussenzweig, M. C. Chromosome translocation, B cell lymphoma, and activation-induced cytidine deaminase. *Annu Rev Pathol* **8**, 79-103, doi:10.1146/annurev-pathol-020712-164004 (2013).
22 Schmidt, F. *et al.* Measuring SARS-CoV-2 neutralizing antibody activity using pseudotyped and chimeric viruses. *J Exp Med* **217**, doi:10.1084/jem.20201181 (2020).
23 Leem, J., Dunbar, J., Georges, G., Shi, J. & Deane, C. M. ABodyBuilder: Automated antibody structure prediction with data-driven accuracy estimation. *MAbs* **8**, 1259-1268, doi:10.1080/19420862.2016.1205773 (2016).
